# An evolutionary approach to identify mammalian adaptive mutations in the avian influenza polymerase complex

**DOI:** 10.1101/2025.10.27.684835

**Authors:** Fernando Capelastegui, Vidhi Dholakia, Damien C. Tully, Daniel H. Goldhill

**Affiliations:** Department of Pathobiology and Population Sciences, Royal Veterinary College, Hatfield, AL9 7TA, UK; Department of Infectious Disease Epidemiology, London School of Hygiene & Tropical Medicine, London, WC1E 7HT, UK; Centre for Mathematical Modelling of Infectious Diseases, London School of Hygiene & Tropical Medicine, London, WC1E 7HT, UK

## Abstract

Avian influenza viruses (AIVs) are a global public health risk; human infection is typically associated with high mortality. While the relationship between several mammalian adaptive mutations and host factors have been described, it is unknown whether additional uncharacterised mutations lead to adaptation. Here, we combine phylogenetic analysis and complementary experimental methods to quantify the impact of novel mutations that emerge at the avian-mammal interface. We constructed phylogenetic trees of mammalian and avian influenza sequences for the polymerase (PA, PB1, PB2) and nucleoprotein (NP) segments and identified potential avian to mammal spillover events. We found >6500 mutations across the polymerase and NP, including known signatures of mammalian adaptation such as PB2 E627K and D701N which occurred independently in mammals 143 and 56 times respectively. We selected 95 mutations which were mostly undescribed and emerged independently multiple times in a range of species and subtypes. Using a minigenome assay in an avian H5N1 backbone to measure the effect of these mutations in human cells we identified PA P28S, NP I425V and G485R as novel mutations leading to polymerase adaptation. In addition, to determine the mechanism of adaptive mutations, we measured polymerase activity in cells lacking a key host factor, ANP32, and cells overexpressing host restriction factors MxA and BTN3A3. Our combined approach revealed novel mammalian adaptive mutations and demonstrated the benefit of combining phylogenetic and molecular approaches in validating novel adaptive mutations.

## Introduction

Influenza A viruses (IAVs) infect and cause severe disease in a range of hosts including humans[1]. Most IAVs circulate in waterfowl, their primary reservoir hosts[2], where they are commonly referred to as avian influenza viruses (AIVs). AIVs are highly adapted to birds, though occasionally spill over into mammals, including humans, often causing disease and high mortality[2,3]. AIVs also pose a global public health risk should they adapt to mammalian and human hosts or reassort with other IAVs, which could lead to the emergence of a novel strain with pandemic potential. Since 2020, highly pathogenic (HPAI) H5 AIVs of clade 2.3.4.4b have been circulating at unprecedented levels year-round in wild birds and spilling-over into poultry where they have had devastating effects[3]. In the US alone, >170 million commercial birds have been culled since 2022[4]. Clade 2.3.4.4b AIVs have also frequently spilled-over into mammalian species causing mass die-offs in marine mammals[5], cats[6,7] and farmed mink[8]. Most recently, there have been three independent emergences and spread among dairy cattle in the USA[3,5,9,10]. Associated human cases, of this clade, and of other AIVs such as H5N1 clade 2.3.2.1e/c, H5N6 and H9N2 continue to fuel concern that AIVs may adapt or reassort to readily infect humans[11,12].

Emergence of a pandemic adapted strain requires a series of mutations across the AIV genome. One key area of interest is the RNA dependent RNA polymerase (made up of PA, PB1 and PB2 segments). A functional polymerase is essential, performing both transcription (vRNA → mRNA) and replication of the viral genome (vRNA ↔ cRNA)[13]. Some polymerase mutations and host factor adaptations have been well characterised and are predominantly driven by adaptations supporting interactions with the essential co-factor Acidic Nuclear Phosphoprotein 32 (ANP32)[14]. Crucially ANP32 proteins act as a strong species barrier in restricting AIV polymerase function in mammalian hosts[14–16]. Due to an exon duplication event, avian ANP32A contains an extra 33 amino acid insertion between the N-terminal leucine-rich repeat (LRR) and the C-terminal low-complexity amino acid region (LCAR)[14,17]. The avian ANP32A LCAR region and the 33 amino acid insertion interact in a negatively charged groove on the polymerase asymmetric dimer complex formed by PB2 627 domains[14,18]. Mammalian ANP32s lack this insertion, which contains residues with mixed charges, leaving the negatively charged LCAR, which cannot interact with the negatively charged polymerase 627 domain without changes that support ANP32 protein interactions[14,18]. PB2 E627K is the most well defined adaptation which biochemically induces the necessary charge change[14,16,19] for ANP32 binding, though other mutations such as Q591R and D701N also support ANP32 binding and stabilisation of the polymerase asymmetric dimer replication platform[16,18,19]. Another protein of interest in host adaptation is the nucleoprotein (NP). NP is closely associated with the polymerase, supporting replication and packaging of the genome[20]. NP also interacts with cellular host factors[20], and in particular, mutations in NP have been shown to be important in overcoming host restriction factors MxA[21,22] and BTN3A3[23]. Previously characterised mutations which interact with MxA and BTN3A3 to support evasion and include Y52N[24], 100I/V,[24] 283P[24] and 313Y/V[23] respectively.

The extent to which there are additional mutations which adapt the AIV polymerase and NP to mammalian cells and host factors is unknown. Previous experimental approaches to identify and characterise mammalian adaptive mutations have provided direct biological evidence of adaptation, though experimental evolution in model organisms e.g. ferrets is costly, constrained by host-specific biology, and subject to ethical and gain of function considerations[25–27]. Recent advances in molecular techniques such as deep mutational scanning (DMS) have provided an alternative in the form of high-throughput in vitro cell-culture systems that allow precise mapping of viral genotype-phenotype relationships. DMS selects on a comprehensive library of all possible mutations at scale[28] and has been used to characterise the effects of mutations in a range of IAV phenotypes including polymerase activity[22,29–34]. However, DMS can be resource-intensive limiting its accessibility and while comprehensive, mutants are not always linked back to real-world emergences constraining its broader application.

Bioinformatic and comparative genomic approaches have also been used to identify mammalian host markers[35,36] from existing sequence data as they are less resource intensive than experimental methods and can be combined with epidemiological and phylogenetic analyses to infer biological and evolutionary contexts. Recent interest in machine learning techniques also promises innovative data-driven approaches to predict zoonotic transmission risk and pathogenicity from annotated sequence data[37–39]. However, these approaches can be susceptible to biases in the sequences used, over-influenced by key mutations such as PB2 T271A or E627K and have limited capacity to resolve the effect of individual mutations[39]. Crucially, findings from bioinformatic approaches are rarely experimentally validated which is necessary to confirm adaptation. However, some studies have combined computational and experimental methods by analysing human isolates of H5N1[40,41] and H7N9[42], identifying mutations of interest, and validating polymerase mutations using minigenome assays in human cells. These studies have looked at mutations in a small number of spillovers, but despite 1000s of mammals infected with AIVs having been sequenced, there have yet to be any large-scale phylogenetic analyses to identify other potentially adaptive mutations.

Here, we combine phylogenetic analyses to identify mutations arising from real-world spillover events with minigenome assays to validate their effect on the polymerase (Figure 1). Through this novel integrative approach, we leveraged publicly available IAV sequences and constructed large phylogenetic trees for each segment of the polymerase and NP. We identified independent mammalian spillovers and collated thousands of associated mutations. Using a minigenome system with an avian H5N1 backbone, we validated the impact of 95 selected mutations on polymerase activity and characterised their mechanism of action. Our methodology revealed large scale diversity in adaptation across the viral polymerase.

**Figure 1.**
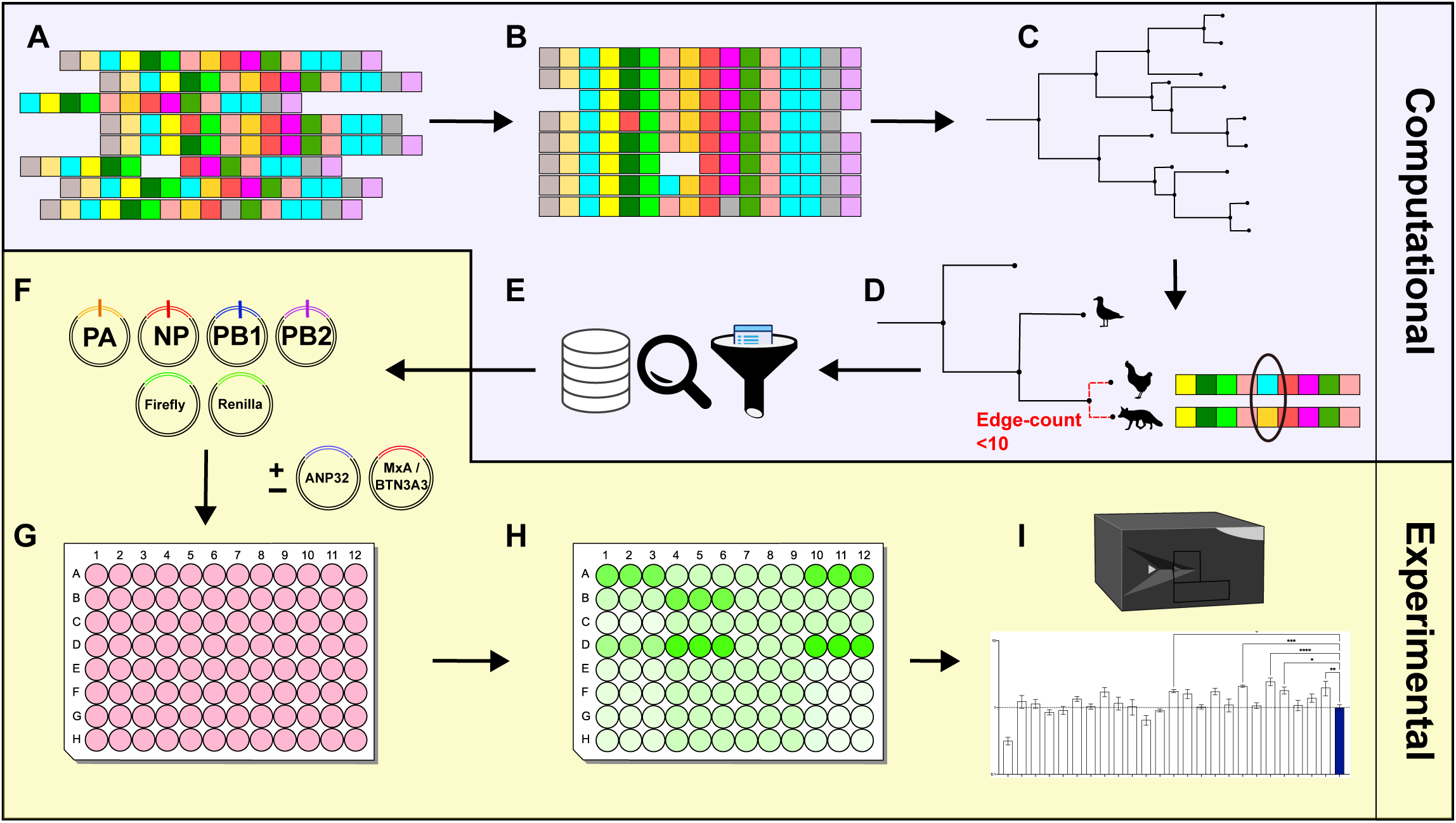
Schematic of computational and experimental pipeline. For more detailed methods refer to methods section. Briefly; **(A)** Avian and mammalian sequences (excl. human H1N1 and H3N2) were downloaded for each segment of the polymerase (PA, PB1, PB2) and NP. **(B)** Sequences were then processed, cleaned, and aligned in MAFFT[44] (v7.409)**. (C)** IQTree2[45] was used to infer maximum likelihood phylogenetic trees for each segment. **(D)** A distance matrix of edge counts for each tip was created; mammalian tips were paired with their closest avian tip. Only pairs within 10 edge counts were kept. Pairwise comparison revealed mutations between selected pairs; pairs with more than 20 amino acid differences were discarded. **(E)** Mutations were tabulated and filtered to remove ambiguous residues. Additional metrics were calculated to infer independent spillover, species, subtype and amino acid frequency at residues across host type. **(F)** A selection of mutations were selected for experimental analysis. Individual mutations were introduced into a H5N1 minigenome backbone by PCR site directed mutagenesis and transfected into HEK 293T cells **(G).** NP minigenomes were run with restriction factors MxA and BTN3A3 overexpressed. ANP32 specific experiments were carried out in ANP32A/B/E triple knockout eHAP cells with ANP32A+B exogenously expressed. **(H)** Minigenome luciferase results were read and quantified **(I).** Schematic illustration made in InkScape V1.4.2[81] using symbols and silhouettes from PhyloPic.com and bioicons.com.

## Results

To identify possible mammalian adaptive mutations, we retrieved ∼57,000 sequences for each polymerase segment and NP across all IAVs uploaded to GISAID[43] (Table S1). We excluded human H1N1 and H3N2 isolates as they would be unlikely to reveal novel mammalian mutations emerging zoonotically whilst adding significant amounts of unnecessary computation. Briefly, sequences were downloaded in batches based on their uploaded metadata into avian, mammal and human categories. The sequences were combined by segment, processed and aligned using MAFFT software[44]. Using IQTree2[45], we constructed maximum likelihood phylogenetic trees using the automatic model finder (Table S1).

### Mutations emerge independently and frequently across the whole polymerase

Next, we analysed the topology of each phylogenetic tree and created a distance matrix based on edge counts for every tip. From this we identified the closest avian sequence for each mammalian sequence, identifying 66,031 mammal-avian pairs across all segments. We kept those that were within 10 edge counts of each other and differed by <20 amino acids. This filtering step reduced the total number of pairs down to 7,351, made up of 4,922 unique avian relatives (Table S1). This filtering step maximised capturing likely zoonotic spillover events and mitigated for noise created by unlikely pairings e.g. sequences which are not temporally or genomically linked, other than by long branches on the tree. Low sampling coverage in rarer subtypes could often lead to long branches and spurious relationships. This was often seen in swine/avian pairings due to a paucity of avian sequences compared to swine. By comparing the sequences of selected pairs, we conservatively identified 1,825 mutations in PB1, 1,822 mutations in PB2, 1,883 mutations in PA and 1,059 mutations in NP. We found the highest number of mutations emerged in PA despite being the smallest of the three polymerase segments, followed by PB2 and PB1 (Figure 2A). Unique emergences and host species were both highest for PB2 followed by PA and PB1 (Figure 2B). This is somewhat expected as many PB2 and PA mutations have been linked to host adaptation and the function of PB1 may be more tightly constrained[46]. *Complete data and interactive figures can be found in supplementary figures S1-4 and interactive supplementary HTML files 1-3*.

**Figure 2.**
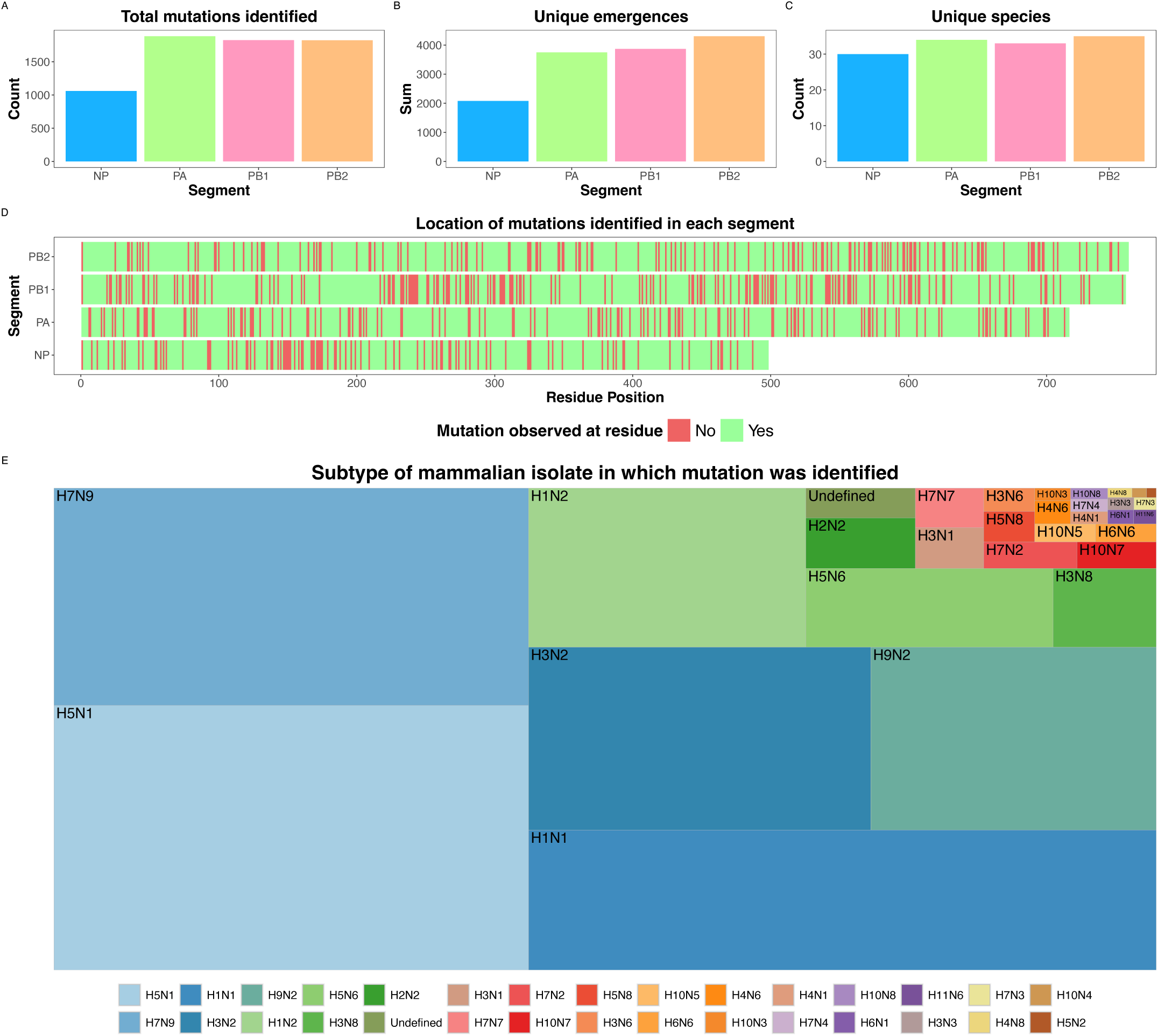
Overview of bioinformatic data analysis divided by unique polymerase segment (PA, PB1, PB2) and NP. **A**: Total sum of unique mutations identified from suitable pairs of avian and mammal sequences. **B**: Total sum of unique emergences of all mutations as defined by how many time a mutation was identified with a unique avian per mutation-identifying avian-mammal pair. **C:** Count of unique host species (determined by GISAID sequence ID metadata **D:** Location of residues across the length of each protein segment with at least one mutation identified. **E:** Distribution of subtypes based on mammalian isolate from which mutations were identified

We calculated the number of independent emergences for each mutation and found mammalian mutations emerged independently a total of 3,865 times in PB1, 4,300 in PB2, 3,745 in PA, and 2,079 times in NP (Figure 2B). Mutations were detected in over 30 mammalian hosts (NP=30, PA=34, PB1=32, PB2=35) (Figure 2C). The most common species were humans and swine, though there were also frequent spillovers to foxes, mink, canine and equine hosts. Mutations were identified in ∼80% of residues throughout each segment. Consistent with earlier results, we observed a higher proportion of mutations across the length of PA (81.7%), PB2 (80.5%) and NP (79.9%) than PB1 (76.9%) (Figure 2D). We detected mutations in mammalian sequences from at least 30 subtypes (Figure 2E). The most common subtypes were H5N1, H7N9, H1N1, H3N2 and H9N2 which aligns with reported AIV spillovers and extensive surveillance of swine populations[11,47].

### Known adaptive and novel mutations frequently emerge

To validate our approach we cross-checked our data against known adaptive mutations reported in the literature. As expected, we observed a high frequency of well characterised mammalian adaptive mutations such as PB2 E627K, D701N and Q591K emerging independently (143, 56 and 22 times respectively) (Figure 3) in a number of different species (13, 14 and 6) and subtypes (10,12 and 5) (Figure S3 and supplementary HTML 1). Other known mammalian adaptive mutations also scored well in our analysis. For example NP V353I, and R351K, both associated with MxA resistance[21], emerged independently a number of times (10 and 11 times respectively). These mutations occurred in a range of hosts including humans, swine, canines and felines emphasising them as broadly mammalian adaptative.

**Figure 3.**
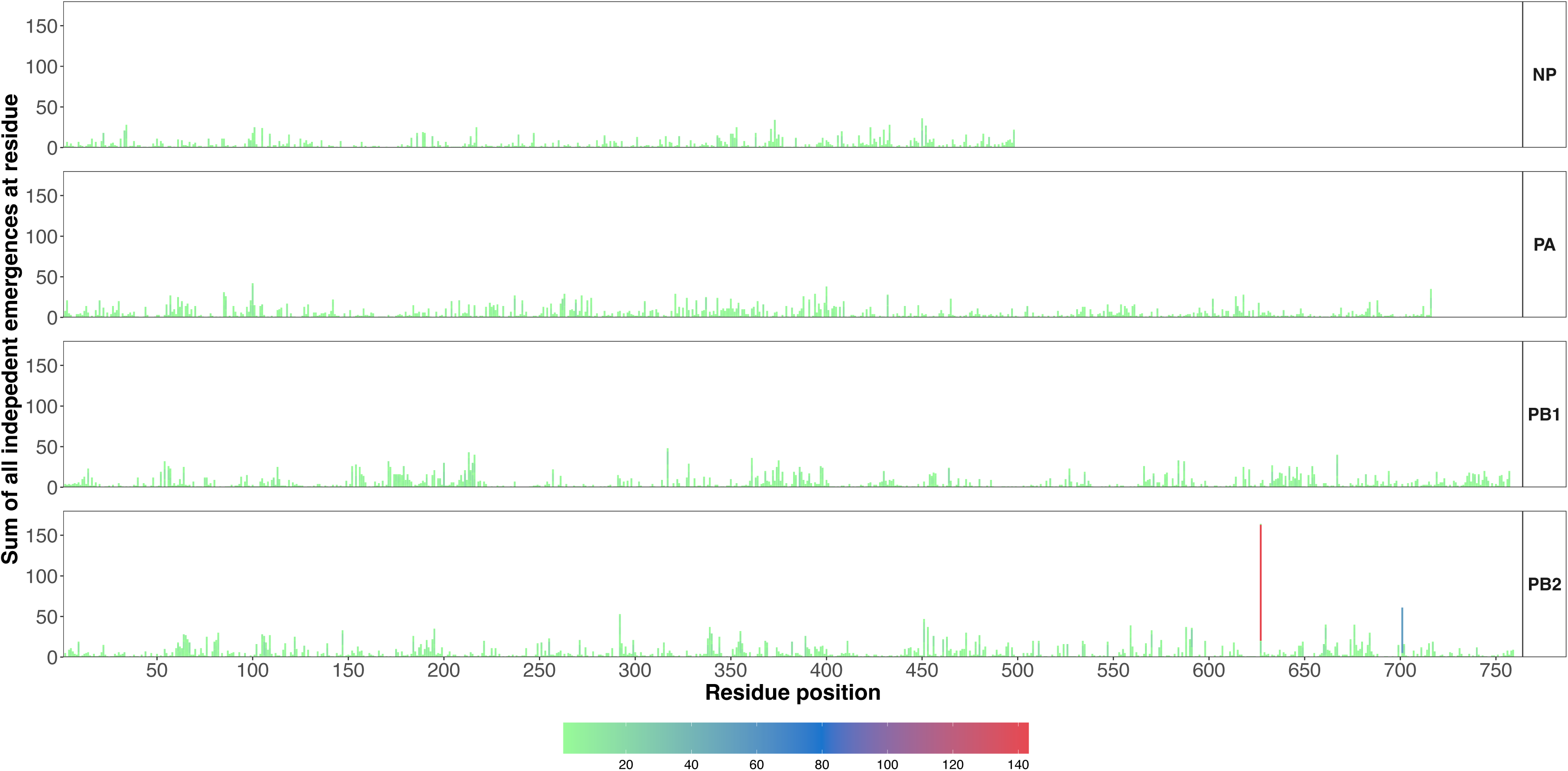
Sum of independent emergences at residue stacked by individual mutation for each polymerase segment and NP. Independent emergence classified as a unique avian sequence for each mammal-avian pair where mutation was identified.

Next, we investigated novel sites with large numbers of mutations. Unlike known adaptive sites such as PB2 627, we noted that some sites showed a large diversity of mutations without a dominant mutation or with bidirectional mutations. For example, at PB2 292, there were 151 changes split across 13 different mutations. The most common mutations were M292I and T292I, yet the reverse mutations were seen relatively frequently, with few independent emergences. Thus, it is necessary to distinguish mammalian adaptations from frequently occurring mutations at variable sites by considering mutation directionality, residue frequencies, species and subtype rather than solely using mutation counts.

To select mutations for further characterization, we ranked mutations by frequency, independent emergences, species and subtype diversity. By sorting on these metrics, we heuristically triangulated mutations and, by simultaneously searching for them in the literature, we found many mutations which provided interesting signals but were yet to be characterised. We subsequently selected 95 mutations of interest across all four segments for experimental characterisation based on these measures. Details can be found in Table S2 which contains mutation specific attributes and reasons for inclusion. For example, P28S in PA occurred in 34 avian-mammal pairings (of which 6 times independently) in humans and swine across 4 subtypes; H1N2, H3N2, H1N1, H7N9. In PA, residue 28 serine (S) amino acid frequency is also much higher in mammals, (26.9% vs 0.1%) which also led to its inclusion, though this is likely due in part to over-representation of swine isolates. In NP, we saw that V33I arose 33 times, 10 of which independently, in a range of species; swine, humans, sea lions, canines, and a range of subtypes; H1N1, H1N2, H5N1, H7N9, H9N2, H3N2. We observed that the residue isoleucine (I) was much more common in mammals than birds at residue 33 (65.21% vs 5.14%) highlighting this as a potential, undescribed mammalian adaptive mutation.

### Selected mutations aDect polymerase activity and interact with ANP32 proteins

To test whether the selected mutations adapted the polymerase to mammalian cells, we introduced individual mutations into expression plasmids containing a WT avian H5N1 polymerase backbone (A/turkey/England/50-92/1991(H5N1)). Using a minigenome experimental system, we quantified polymerase activity in human 293T cells for each mutation and normalised these results to the WT backbone (Figure 4). PB2 E627K was included as a positive control and gave high activity (21x increase on WT, p<0.0001). In addition to E627K, we saw increased activity from well characterised mutations PB2 D701N, T271A, as well as some effect from PB2; S286G[42], Q591R, PB1; L598P, N105S[48] and P708S[49] which have previously been characterised. Of note, PB2 S286G, PB1 L598P and P708S boosted activity significantly 10.9x, 13.1x and 3.9x respectively compared to WT (p<0.0001, p<0.0001, p=0.0103). In PA we also confirmed the activity of several known mutations; T85I[50,51], K356R[52], K362R and K497R[53] (Figure 4). Of the PA mutations we found T85I had the greatest effect, increasing activity by 4.4x compared to WT (p<0.0001). There were also some novel mutations which increased polymerase activity to varying extents but have not been previously characterised in an AIV H5N1 backbone such as PA P28S, PB1 S642N, and P708S.

**Figure 4.**
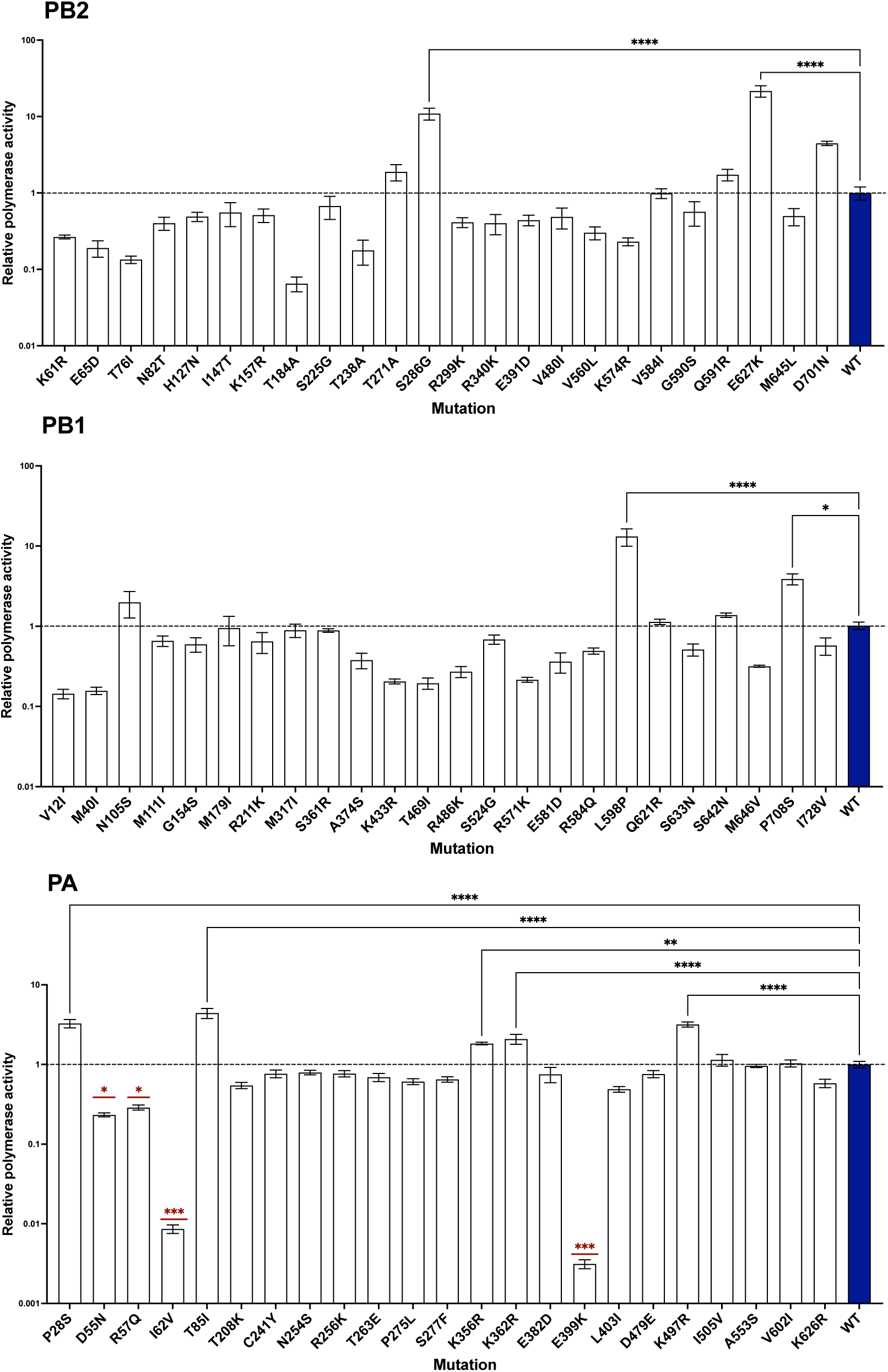
Minigenome assay of polymerase segments (PB2, PB1, PA) with selected mutations in human 293T cells. Polymerase activity displayed as Firefly/Renilla normalised to wild type, with SEM bars. Results of one-way ANOVA comparing WT to mutant are displayed. (*P ≤ 0.05, ** P ≤ 0.01, *** P ≤ 0.001, **** P ≤ 0.0001). Combined results from 2 experimental repeats are shown.

We next wanted to determine if the mutations that significantly boosted polymerase activity were specifically adapting the polymerase to co-opt mammalian ANP32 proteins. We selected mutations from PB2, PB1 and PA which had significantly improved polymerase activity, and repeated minigenomes in human ANP32A/B/E triple knock out eHAP cells (eHAP TKO) with either chANP32A or huANP32A+B exogenously expressed (Figure 5). We found that generally, the mutant polymerases were still very much avian adapted, performing much better in the presence of chANP32A (Figure 5A). When normalised to their respective treated WT, as expected we found PB2 E627K significantly boosted polymerase activity when huANP32A+B were present vs WT (36.2x increase on WT p<0.0001) and further appeared to be deleterious to polymerase function when chANP32A is present (0.4x of WT activity). Relative to their respective WT controls (Figure 5B), PB2 S286G and PB1 L598P both boosted polymerase function in the presence of both huANP32A+B (17.5x; p<0.0001 and 5.6x; p=0.0190 respectively) and chANP32A (5.3x; p=0.0374 and 2.7x; p=NS respectively), though more so for huANP32A/B suggesting they could be specifically adapting the AIV polymerase to mammalian ANP32A/B. Although PB2 S286G and PB1 L598P have previously been characterised as mammalian adaptive mutations[54], this confirms the mechanism of action linking them to mammalian ANP32 proteins. The remainder of the mutations tested in this assay boosted polymerase activity relative to each WT regardless of which host ANP32 proteins were present. This suggests that the mechanism of adaptation for these mutations is not directly linked to specificity for mammalian ANP32.

**Figure 5.**
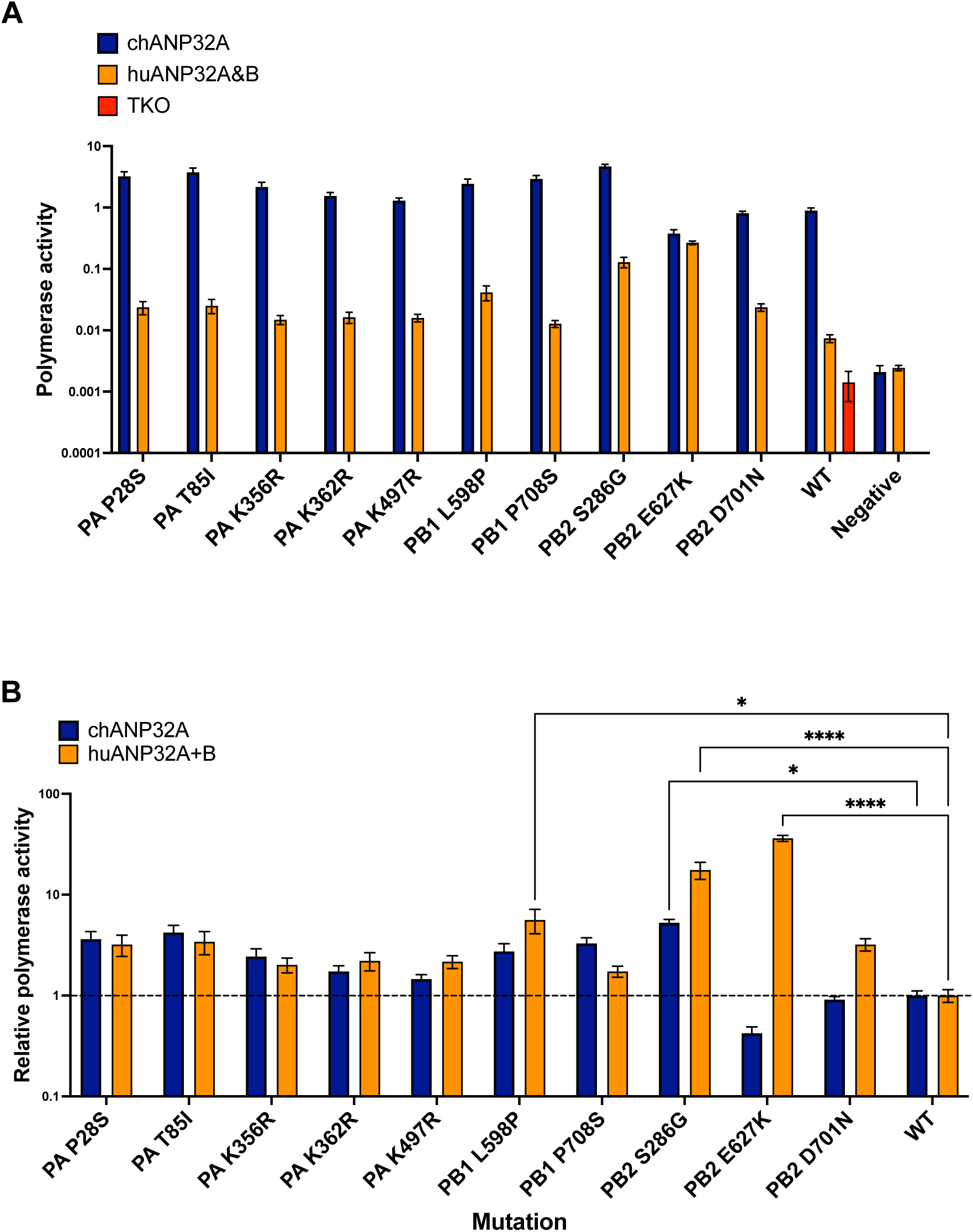
Minigenome assay of select polymerase mutations in ANP32A/B/C triple knockout human EHAP cells. Either human ANP32A+B (huANP32A+B) or chicken ANP32A (chANP32A) was exogenously expressed. Results of two-way ANOVA comparing WT to mutant for each treatment group are displayed. (*P ≤ 0.05, ** P ≤ 0.01, *** P ≤ 0.001, **** P ≤ 0.0001). Combined results from 2 experimental repeats are shown. **A:** Polymerase activity displayed as raw Firefly/Renilla readings, with SEM bars. WT in TKO cells displayed for reference. Wild type negative control and treatment negative control displayed for comparison. **B**: Relative polymerase activity displayed normalised to wild type for each treatment group, with SEM bars.

We found that most novel mutations we selected from the bioinformatic pipeline did not increase polymerase activity in this background compared to WT, nor did any mutations increase polymerase activity to the level of E627K. We found that some mutations significantly reduced polymerase activity to a greater extent. Notable examples were PA E399K and I62V where polymerase activity was 0.0031x and 0.0086x that of WT (p=0.0006 and p=0.0005 respectively). Although not statistically significant, similar examples of mutations that reduced polymerase activity were found across PB1; V12I, M40I and in PB2; T76I and T184A.

### Nucleoprotein mutations can overcome host restriction and boost polymerase activity

To assess the role of NP in mammalian adaptation, we ran a minigenome of selected mutants as above in 293T cells (Figure 6A). NP F313V, which is a known BTN3A3 resistance mutation, showed a significant increase (1.7x, p=0.0497) in polymerase activity. Two other mutations previously described in the literature, T350K and Q357K showed a boost to polymerase activity though it was not statistically significant in this assay. Several novel uncharacterised mutations I425V, M426L and G485R were shown to boost polymerase activity by 2.4x, 1.8x and 2x respectively (Figure 6A). As NP mutations have previously been shown to be important in evading host restriction, we repeated minigenomes with human restriction factors MxA and BTN3A3 overexpressed. We detected signals of MxA and BTN3A3 evasion in several known and novel mutations (Figure S5). Informed by this data we selected several mutations (V100I, L283P, F313V, T350K, Q357K, R400K, I425V, M426L, G485R) for further confirmation in repeat side-by-side experimental comparisons at baseline conditions and with either MxA or BTN3A3 overexpressed. In the presence of restriction factor overexpression, WT polymerase activity was impaired. We found polymerase activity was knocked down in this backbone more effectively using MxA compared to BTN3A3 (Figure 6B). All selected mutations showed increased polymerase activity compared to WT when we overexpressed MxA and BTN3A3 (Figure 6B). By normalising mutant polymerase activity to their respective WT within each treatment group, we confirmed that known mutation L283P boosted activity to a greater extent when either MxA or BTN3A3 were overexpressed compared to WT at baseline (5.4x and 6.3x vs 3.1x increases compared to respective treated WT). We also confirm F313V can overcome BTN3A3. When BTN3A3 was overexpressed F313V activity was 4.6x higher (p<0.001) compared to WT, but only 2.3x and 2x higher nor significantly increasing polymerase activity at baseline or when MxA was overexpressed (Figure 6C). We found that G485R significantly boosted polymerase activity when MxA was overexpressed (4.02x, p=0.022) and in baseline conditions (3.88x, p=0.033). Albeit not statistically significant, G485R also increased activity under BTN3A3 overexpression conditions compared to its respective WT (3.39x, p=0.114). Although I425V also boosted polymerase activity when either MxA and BTN3A3 were overexpressed, it only significantly increased polymerase activity at baseline (5.2x, p=0.004) (Figure 6C). The mechanism of adaptation for these novel mutations is therefore unclear as they generally increased polymerase activity proportionally both at baseline and when restriction factors were overexpressed.

**Figure 6.**
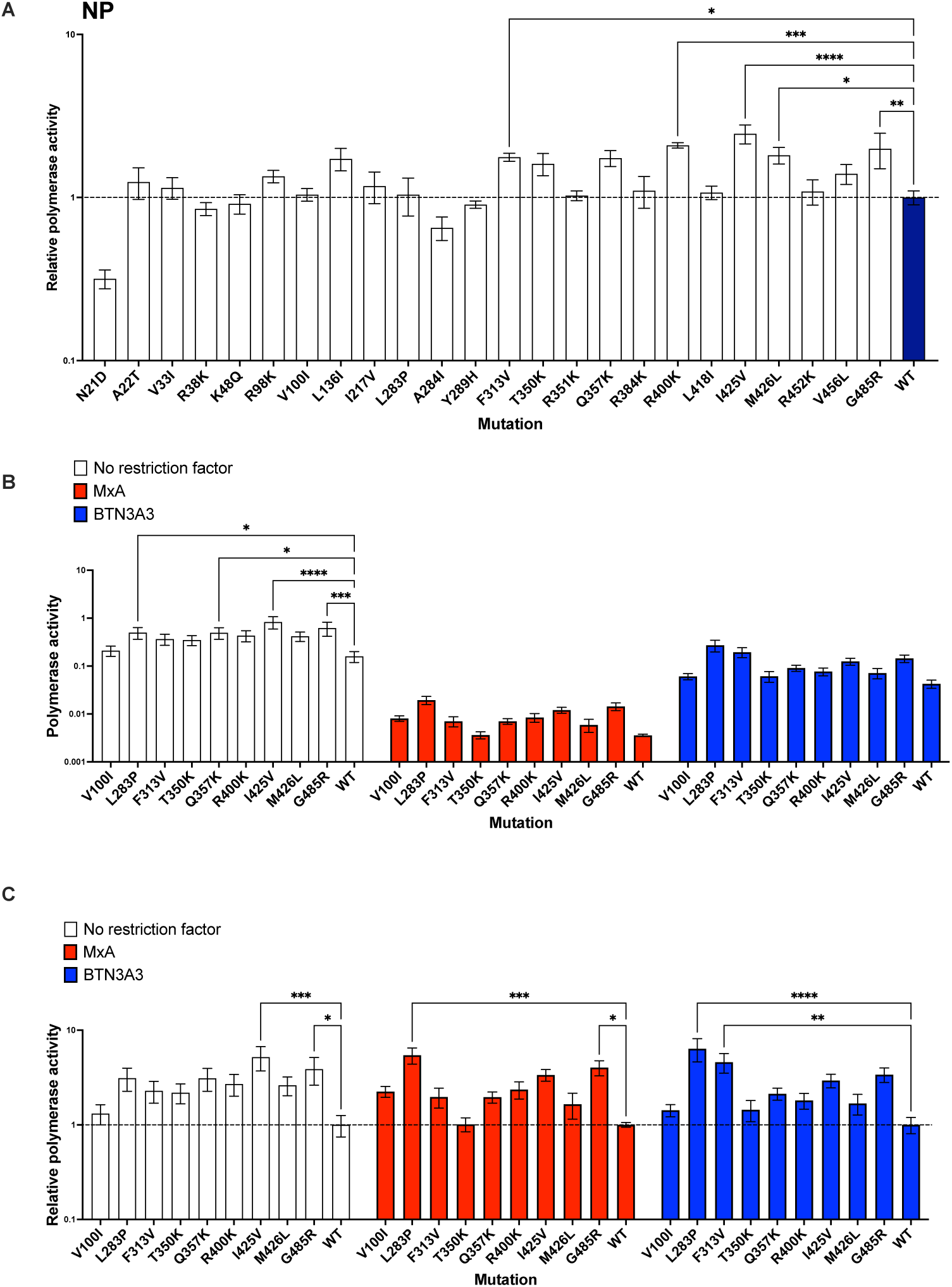
**A:** Minigenome assay of NP mutations in human 293T cells. Polymerase activity displayed as Firefly/Renilla normalised to wild type, with SEM bars. Results of one-way ANOVA comparing WT to mutant where p value greater or equal to 0.05 displayed. (*P ≤ 0.05, ** P ≤ 0.01, *** P ≤ 0.001, **** P ≤ 0.0001). Combined results from 2 experimental repeats are shown. **B:** Raw polymerase activity readings for select NP mutations in human 293T cells with restriction factor BTN3A3 or Mx1 exogenously expressed. Polymerase activity displayed as Firefly/Renilla with SEM bars. Results of two-way ANOVA comparing WT to mutant by treatment group are displayed. (*P ≤ 0.05, ** P ≤ 0.01, *** P ≤ 0.001, **** P ≤ 0.0001). Combined results from 2 experimental repeats are shown. **C:** As in panel B, with polymerase activity normalised to respective wild type per treatment.

## Discussion

Through an evolutionary bioinformatics screening approach, we have identified thousands of potentially mammalian adaptive mutations across the polymerase and NP genes. By comparing closely related sequences across the entire IAV phylogenetic landscape, we detected mutations arising repeatedly in biologically relevant avian-mammal transitions at scale. This novel resource is an inventory of likely mammalian adaptive mutations, revealing extensive diversity and possible combinations of mutations spanning ∼80% of the polymerase structure. As expected, key mammalian adaptive markers such as PB2 E627K and D701N occurred more frequently with more independent occurrences across a broader range of species and subtypes. Many other mutations showed signals consistent with mammalian adaptation yet remain uncharacterised. By mapping the top 50 most frequent mutations onto relevant structures (Figure 7, HTML 2&3), we highlight that diverse areas of the polymerase contribute to mammalian adaptation without obvious hotspots other than the 627 domain. We identified novel adaptive mutations NP I425V, G485R and PA P28S but we highlight that bioinformatically identified mutations should not be assumed to be mammalian adaptive as we found that most mutations showed no increase in polymerase activity. This corresponds with other studies which have combined experimental and bioinformatic approaches[41,55] and further points to unresolved mechanisms governing polymerase adaptation.

**Figure 7.**
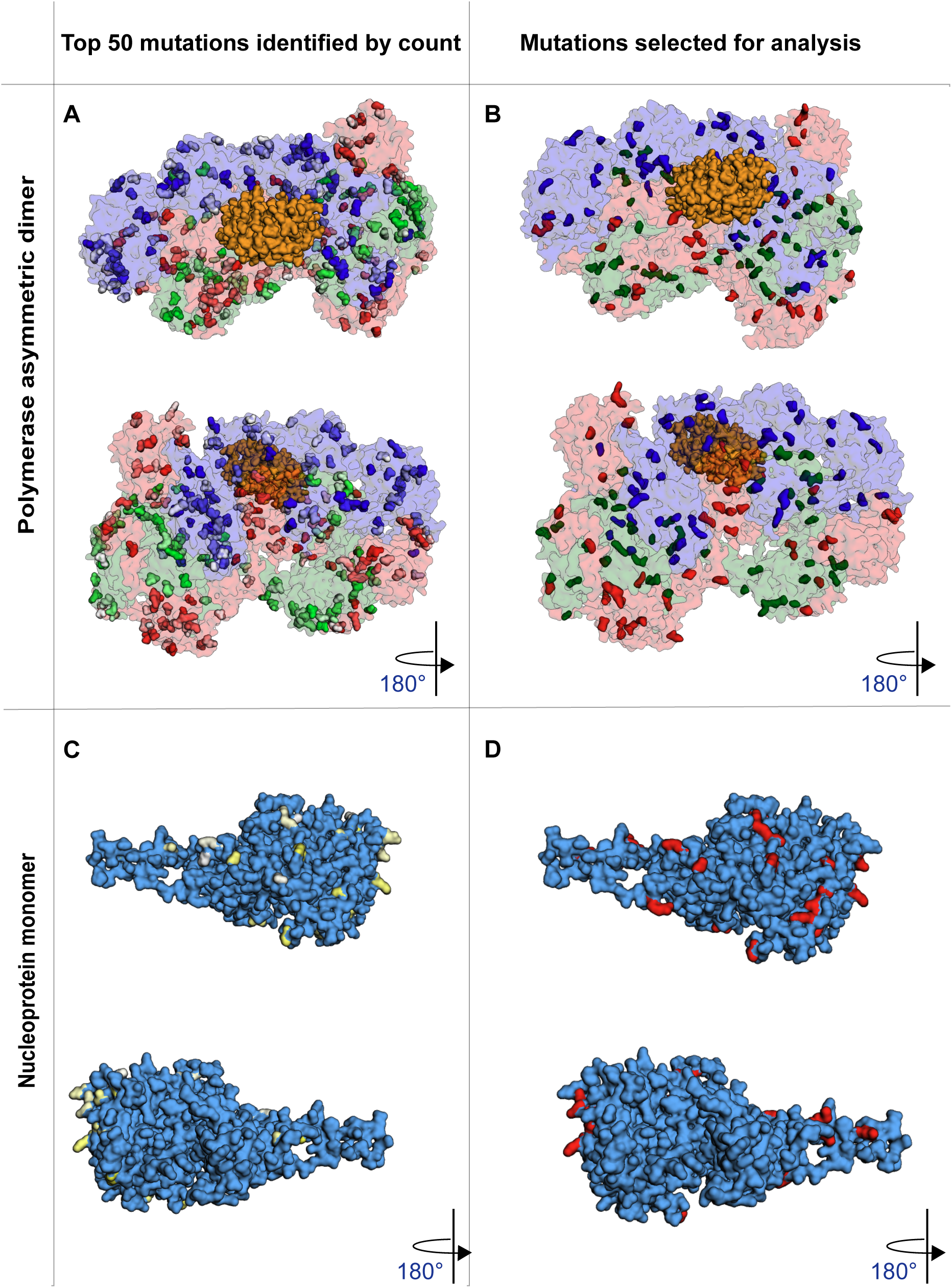
H5N1 asymmetric polymerase dimer bound to human ANP32B split by top 50 mutations observed by count **(A)** and those mutations selected **(B)**. (PDB: 8R1J. Red: PA, Blue: PB2, Green: PB1, Orange: ANP32B). NP H5N1 monomer (**C & D**) as above). PDB:2Q06

In addition to characterising whether mutations were broadly mammalian adaptive, we also investigated whether the mechanism behind our adaptive mutations involved the well characterised host co-factor ANP32. We demonstrated that PB1 L598P and PB2 S286G support improved use of human ANP32A and ANP32B proteins. These mutations have previously been associated with human viruses[42,54] and PB1 L598P has been shown to compensate for the lack of PB2 E627K in an H5N1 backbone[54] suggesting an ANP32 interaction. However, the molecular mechanism remains unclear with neither mutation being located in direct proximity to the ANP32 binding region in the asymmetric dimer complex, nor in the symmetric dimer interface[19,56] (Figure S6). These mutations may be additive when in combination with other changes, or potentially biasing the polymerase towards asymmetric dimerization which has been linked to interactions with suboptimal mammalian ANP32 proteins[42,57].

In PA, we identified the novel mutation P28S which occurred in swine and humans and more commonly in mammalian than avian sequences. PA P28S showed no increase in polymerase activity with human ANP32 over avian ANP32 and therefore the mechanism behind mammalian adaptation remains unknown, though as PA P28S is located in the N-terminal endonuclease domain and not in any polymerase dimer interface, direct ANP32 interactions were unlikely (Figure S6). While we did not set out to identify mutations in PA-X, it has been shown that Proline (P) at position 28 vs a Leucine (L) is an important determinant of PA-X shut off and polymerase activity when comparing a A/California/04/2009 (H1N1pdm09) backbone to A/WSN/33 (H1N1)[58]. As we mainly identified P28S in swine isolates it would be interesting to further investigate this effect and mechanism in a range of backgrounds and experimental systems as the effects of PA-X activity are lineage and host-specific[58,59].

Our results have expanded the number of NP mutations linked to increased polymerase activity and overcoming host immune evasion. Previous evidence has shown that individual NP mutations including L283P, F313V and V100I can overcome the restriction factors MxA[60] and BTN3A3[23]. We showed that novel mutations I425V and G485R boosted polymerase activity despite both restriction factors being overexpressed. However, these mutations boosted polymerase activity in all conditions suggesting their main mechanism of adaptation is not directly related to overcoming these host restriction factors. Mutation I425V is found in the tail loop region of NP and has been bioinformatically and experimentally associated as a marker of human adaptation though its function remains unknown[61–63]. 425V was much more common in mammals (62.55% vs 0.35%) and linked to human and swine isolates. Previous in vitro experiments showed I425V did not improve viral growth in a duck H1N1 background^61^, nor particularly increase polymerase activity in a minigenome with a pigeon H7N9 background^62^. Previous work has validated this site amongst others as important in viral antigenic epitope variation which allows escape of recognition by cytotoxic T-lymphocytes[64] though this mechanism would not explain the effects seen in cell culture here. Less is known about NP G485R; we found that G485 is highly conserved in both avian and mammalian isolates, occurring in >99% of sequences whereas 485R is very rare but its frequency is nearly ten times higher in mammals than in birds (0.2% vs. 0.03%). There is limited evidence 485R may support increased transmission experimentally in ferrets in conjunction with PB2 E627K when using fox H5N1[65] or human H7N9[66] isolates, but its effect on polymerase activity had not been tested. Structurally, both 425 and 485 may be important in NP homo-oligomerisation, which is important for stabilisation of viral RNPs and for both transcription and replication of the viral genome (Figure S7). Although our analysis did not investigate the co-occurrence of these mutations, it would be interesting to explore further.

Interestingly our pipeline revealed that G485R emerged independently 12 times, in a range of species; foxes, swine, racoons, humans, seals and mink. This complements recent surveillance efforts of mammals infected with H5N1 2.3.4.4b which reported the emergence of G485R without experimentally validating the mutation as mammalian adaptive[67,68]. We were able to confirm this mutation may be adapting the polymerase to mammalian hosts and show that although it is a rare mutation occurring at low frequencies, it is relevant to our understanding of viral adaptation and future preparedness efforts should this mutation occur more frequently in circulating H5N1 strains or emergent AIVs. Similarly, in the context of the dairy cow outbreak in the US, our analysis was also able to detect signals for key mutations which have emerged. For example, PA K497R which is a known mutation[53,69], and a hallmark of the cattle outbreak in the US[69] stood out from our analysis; R497 occurs more frequently in mammals than avian sequences (23.78% vs 1.03%) and also emerged in a range of subtypes (H5N1, H1N2, H3N2, H1N1 and H7N9).

In the context of pandemic preparedness for “Disease X”, our approach can be applied to a range of possible zoonotic pathogens as a tool for both retrospective and real time characterisation of candidate adaptive mutations. Influenza is somewhat unique having numerous independent spillovers from birds to mammals coupled with extensive sampling and coverage compared to other viruses. By contrast, this methodology is not well suited to SARS-CoV-2, which despite its zoonotic origin, spread predominantly within humans following an initial spillover event[70,71]. Pathogens such as West Nile virus and Nipah virus would be more suitable pathogens to apply our methodology to as there are frequent spillovers into humans[72,73] and host populations are clearly defined. The continued expansion of global sequencing coverage coupled to open access databases e.g. Pathoplexus[74] will further reduce barriers to apply this approach.

### Limitations

A major limitation in this study is the use of a single H5N1 avian backbone to experimentally characterise mutations of interest. This backbone may not be appropriate to show adaptation for mutations that we identified emerging in a diverse range of subtypes and genetic backgrounds. Furthermore, we also only investigated one mutation at a time, which does not reflect epistatic interactions with other mutations emerging at the same time, or present in the specific genetic background. Together, these factors may explain why not all of the mutations we tested increased polymerase activity. Here, we were also only able to investigate <100 mutations but our bioinformatic screen highlighted many promising mutations that remain untested such as PB2 N102S which occurred 8 times independently in a range of species, or PA V100I which emerge 13 times independently. A logical next step is to expand the panel and measure the effect of multiple mutations across the same and different polymerase segments.

While minigenome assays in 293T human cells are a quick tool to assess polymerase activity, they do not perfectly reflect biological reality. These minigenomes only quantified mRNA production using luciferase as a proxy, which may have masked other aspects of polymerase activity and broader viral fitness e.g. viral packaging or immune response. It is possible that the mechanism of adaptation for these mutations is species specific and a biological effect may not be visible in human cells. However, the emergence in multiple species of the mutations we tested is strong evidence for mammalian rather than species specific adaptation. Finally, we acknowledge that there will be large biases in data completeness and sequencing coverage across species and regions when using publicly available sequences can influence phylogenetic tree completeness and topology. Our estimates are therefore conservative and the true number of spillover events is likely underestimated due to sequencing gaps.

### Conclusion

Our method establishes a proof of concept framework that couples evolutionary bioinformatics with complementary experimental methods to validate mammalian adaptive mutations. This strategy could be applied to other IAV proteins such as HA and NA and adapted to zoonotic viruses with recurrent cross-species transmission. Further exploration of the bioinformatic and experimental data presented here will refine our understanding of AIV mammalian adaptation which remains a public health priority.

### Materials & Methods

### Bioinformatics – data processing

GISAID[43] was queried on 04 March 2024. Sequences for every subtype for PB1, PB2, PA and NP segments were downloaded using the filters “Mammal”, “Human” and “Avian” species. Human sequences of subtypes H1N1 and H3N2 were not queried due to volume and the endemic seasonal nature of these viruses. For each segment, raw sequences were deduplicated and aligned using MAFFT[44] software (v7.490) under standard parameters. Only sequences that were majority complete (< 10 AA missing) were kept. Sequences were cleaned and trimmed using a combination of Trimal[75] and CIAlign[76] software. Maximum likelihood (ML) phylogenetic trees were inferred using the IQTree2[45] software. The “AUTO-fast” setting was used due to the size of the alignments. A model was selected by testing various options in the ‘VIRAL’ library. All further analysis was carried out in R[77] unless specified. All scripts and processing parameters can be found in the manuscript GitHub. The alignment files were processed to create a metadata lookup of subtype and species for each sequence. Subtype was determined by extracting the string containing “HxNx” from the GISAID isolate, and the small proportion of sequences where this was not included were assigned the label “Undefined”. Species was also determined from the GISAID isolate name, and where not available, we assigned the label “mammal” or “avian”. Scientific and common names were combined into one label i.e. “Vulpes Vulpes” and “Red Fox” were combined into “Fox” for suitable grouping.

### Bioinformatics – phylogenetic analysis

Each tree was imported into R and linked to sequence species and subtype metadata. Due to the size of each tree, they were split into sub trees for more efficient computation. To ensure all topologies were captured, each tree was split into smaller subtrees in batches of sequential tips, for the following batch sizes: 500, 5000, 10000, 15000.

For each subtree, a matrix of pairwise distances between each tip was calculated by edge counts using the castor R-package[78]. For each mammalian tip, the closest avian ancestor was identified. As multiple trees were used, the avian-mammal pairings were collated and deduplicated, keeping the closest pair if there were more than one mammal-avian paring. In the case of a mammal tip being paired with 2 or more avian tips of the same distance, all equal pairings were retained. An edge-count distance cut-off of <10 was used to filter out mammal-avian pairs that were more distantly related.

Pairwise sequence analysis between selected mammalian and avian sequences identified mismatches i.e. mutations. The mutations and sequence IDs were collated into a database. Pairs > 20 mutations between them were filtered from the final database to minimise spurious mammal-avian pairings.

### Secondary data analysis

To clean and process the final database of mutations, only mutations with non-ambiguous amino acid residue codes were retained; “X” or other ambiguous codes were discarded. For each mutation, the following metrics were calculated; number of times occurred, average distance between pairs, species number, species type, subtype and subtype count. A count of how many times the mutation emerged independently was created by counting how many times the closest avian relative was unique. This accounts for tree topology where an avian relative is closely related to an outbreak of identical mammal sequences, providing a sense of how important a mutation is if it repeatedly emerges independently. The dataset was enriched with percentage occurrence for each residue for avian and mammal sequences calculated from the alignments used to infer ML trees. Mutations were ranked by each of these metrics and scored based on their order i.e. the most common mutation was scored 1 for that metric. Mutations were manually selected for subsequent laboratory investigation based on these rankings (Table S2). Several known mutations were also selected per segment to act as positive controls and for comparison in this background. Full data output and metrics can be found in the complementary interactive HTML files uploaded to the project GitHub: https://github.com/fernandocapelastegui/evolutionary_approach_mammalian_adaptive_AIV_polymerase_mutations

### Molecular biology

#### Plasmids

pcDNA3.1 expression plasmids (CMV enhancer and CMV promoter) were constructed with H5N1 A/Turkey/England/50-92/1991 PA, PB1, PB2 and NP segment inserts (NEBuilder HiFi DNA Assembly Kit). Mutations were introduced by site-directed mutagenesis using KOD Hot Start DNA polymerase PCR kit (Sigma-Aldrich).

Mutagenesis was confirmed by Sanger sequencing (Source Bioscience). For host factor overexpression assays, pCAGGS expression plasmids containing huANP32A, huANP32B, chANP32A, huMxA and huBTN3A3 were used. Reporter plasmids were pCAGGS huPOLI vRNA expression plasmids containing a Firefly luciferase flanked by non-coding viral like sequence and a pCAGGS huPOLII Renilla luciferase reporter.

#### Cells

Human Embryonic Kidney 293T (293T) cells were maintained in Dulbecco’s modified Eagle medium (DMEM, Gibco) supplemented with 1% Penicillin/Streptomycin and 10% fetal bovine serum (FBS, biosera). Engineered ANP32A, ANP32B and ANP32E triple knockout human haploid cells (eHAP TKO) were maintained in Iscove’s modified medium (IMDM, Gibco) supplemented with 1% Penicillin/Streptomycin (Gibco) and 10% FBS. All cells were cultured at 37 °C and 5% CO_2_.

#### Minigenome assay

48 well TC-treated plates were seeded with 293T cells and cultured for 24 hours to reach approximately 70% confluency. The cells were transfected with the following amounts of plasmids: 40 ng PB2, 40 ng PB1, 20 ng PA, 60 ng NP, 40 ng Firefly luciferase, 40 ng Renilla luciferase. A wild type control containing stock plasmids of unmutated A/Turkey/England/50-92/1991 was included in every assay. Positive controls containing PB2 E627K and negative controls lacking the PB2 plasmid were also included in the baseline assays to ensure transfection efficiency but not included in statistical analyses or displayed in results. More detail can be found in section *Statistical analysis*.

To test for ANP32 interactions, eHAP TKO cells were also transfected as above with either 40 ng chANP32A, 40 ng of both huANP32A and huANP32B, or untransfected without any ANP32.

Initial NP host restriction assays testing all NP mutations were carried out as above and transfected with 60 ng of huMxA or 120 ng of huBTN3A3 encoding plasmid. These amounts were determined a priori and deemed sufficient to determine any signal, strongly restricting WT activity by 95% and 85% respectively. Known NP mutations V100I and F313I acted as positive controls in subsequent smaller assays of selected mutations and were further tested with either 120 ng of huMxA or 120 ng of huBTN3A3 encoding plasmids. WT and -VE controls with or without treatment were included to ensure transfection efficiency but only the WT and +treatment results were included in subsequent statistical analyses and results.

Following transfection all cells were cultured at 37 °C and 5% CO_2_. Cells were lysed 24hrs post transfection using passive lysis buffer. Polymerase activity was measured using Promega dual-luciferase reporter kit on a Tecan iControl Infinite 200 Pro plate reader.

For each assay, each mutation was tested in triplicate and experimentally repeated at least twice on different days. Data from both repeats was combined for statistical analysis and visualisation unless otherwise specified. Polymerase activity was calculated as a ratio of Firefly:Renilla.

#### Statistical analysis

Visualisation and statistical analysis was performed in GraphPad Prism 10 for macOS. V10.4.2. A one-way analysis of variance (ANOVA) was performed to compare the relative polymerase activity (normalised to WT average) across mutants to that of WT. This was performed for the minigenomes containing all variants in a segment. Post hoc comparisons were conducted using Dunnett’s multiple comparisons test.

A two-way ANOVA was performed on raw polymerase activity to assess the effects of genotype (WT Vs mutants) and treatment groups for ANP32 minigenomes (huANP32A+B or chANP32A) and restriction factor genomes (MxA or BTN3A3). Post hoc comparisons were conducted using Dunnett’s multiple comparisons test. To asses mutant effect compared to respective treated WT, normalised relative polymerase activity was calculated and a two-way ANOVA was performed with a post-hoc Dunnett’s multiple comparison test.

#### Structural analysis

Mutations were visualised using R3DMol[79] using R coding software and ChimeraX v1.6.1[80]. Structural figures were created in InkScape V1.4.2[81]

## Supporting information

Supplementary HTML files 1-3

## Acknowledgments

We thank all laboratories that uploaded sequences to GISAID without which this study would not have been possible.

Funding for this study was from Royal Society Grant 231225 and Academy of Medical Sciences Springboard Grant 1049 to DHG. FC was supported by a Bloomsbury fellowship as part of the LIDo Doctoral Training Program (BBSRC). pcDNA3.1 plasmids for SDM were kindly donated by the Hutchinson lab as part of The Influenza Virus Toolkit (CVR, Glasgow). huANP32A&B, chANP32A, huMxA and huBTN3A3 encoding expression plasmids as well as eHAP TKO cells were kindly provided by the Barclay lab, Imperial College London.

## Notes

https://github.com/fernandocapelastegui/evolutionary_approach_mammalian_adaptive_AIV_polymerase_mutations

### Competing Interest Statement

The authors have declared no competing interest.

