## Supplementary HTML files 1-3 for "An evolutionary approach to identify mammalian adaptive mutations in the avian influenza polymerase complex": HTML_1.html

#### Metadata

- Mutation = Mutation identified
- Residue = Residue number
- Amino Acid = Amino acid seen in mammalian sequence
- Count = Frequency of residue occurrence
- Freq. % in avian sequences = Proportion of sequences where that
  amino acid occurred in avian sequences in whole alignment
- Freq. % in mammal sequences = Proportion of sequences where that
  amino acid occurred in mammalian sequences in whole alignment
- Independent mutation occurrence = How many times did this mutation
  arise between a mammal and a bird, where the avian sequence is
  unique
- Species count = In how many mammalian species this mutation was
  identified
- Species diversity = List of species mutation occurred in. Note:
  species label is as accurate as the metadata provided and aggregated
  into common naming conventions. For sequences where metadata did not
  contain sufficient information, species was labelled as “mammal”
- Subtype(s) = Subtype of mammalian isolate in which mutation was
  identified
- Subtype(s) n = Number of different subtypes in which mutation was
  identified

### PB2

### PB1

### PA

### NP

### Unique emergence plot

#### Interactive Figure 3. Sum of independent emergences at residue stacked by individual mutation for each polymerase segment and NP. Independent emergence classified as a unique avian sequences for each mammal-avian pair mutation was identified in.

### Mutation frequency plot

#### Interactive Figure S1. Sum of all mutations at residue stacked by individual mutation for each polymerase segment and NP.

### Amino acid frequency plot

#### Interactive Figure S2. Heatmap of amino acid residues seen in mammalian sequences. Coloured by frequency on a log10 scale. Mutations selected for analysis circled.

### Species diversity plot

#### Interactive Figure S3. Stacked count of unique species associated per mutation. Red line overlaid displays unique species at residue position.

### Mutations selected for analysis
