## Supplementary HTML files 1-3 for "An evolutionary approach to identify mammalian adaptive mutations in the avian influenza polymerase complex": HTML_2.html

H5N1 asymmetric polymerase dimer bound to human ANP32B. Red: PA.
Blue: PB2, Green PB1. PDB: 8R1J

- Top 50 mutations observed by count. Colour gradient represents
  number of occurrences; the lighter the hue, the higher the count.
- Mutations which were selected for experimental analysis.

### Results

#### Top 50 mutations

#### Mutations selected for analysis
