## Supplementary HTML files 1-3 for "An evolutionary approach to identify mammalian adaptive mutations in the avian influenza polymerase complex": HTML_3.html

H5N1 Nucleoprotein monomer (PDB:2Q06 - chain A) and H1N1 trimer
structure (PDB:2IQH)

- Top 50 mutations observed by count. Colour gradient represents
  number of occurrences; the lighter the hue, the higher the count.
- Mutations which were selected for experimental analysis.

### Results

#### Monomer - Top 50 mutations (NP)

#### Monomer - Mutations selected for analysis (NP)

#### Trimer - Top 50 mutations (NP)

#### Trimer - Mutations selected for analysis (NP)
