## Supplementary figures for "An evolutionary approach to identify mammalian adaptive mutations in the avian influenza polymerase complex"

**Table S1.** Data processing and metadata

| | **Host** | **PB2** | **PB1** | **PA** | **NP** | **Total** |
|---|---|---|---|---|---|---|
| **Raw data** | Avian | 41,178 | 40,414 | 40,862 | 40,734 | 163,188 |
| | Human | 2,265 | 2,214 | 2,300 | 2,262 | 9,041 |
| | Mammal | 14,184 | 13,886 | 14,392 | 14,544 | 57,006 |
| | All | 57,627 | 56,514 | 57,554 | 57,540 | 229,235 |
| **Cleaned alignments** | All | 54,168 | 53,856 | 54,350 | 47,579 | 209,953 |
| **IQTree2 model** | All | FLAVI+G4 | FLAVI+G4 | FLAVI+G4 | FLAVI+I+G4 | |
| **Mammal-Avian pairings pre cleaning** | | 16,939 | 15,793 | 16,680 | 16,619 | 66,031 |
| **Mammal-Avian post cleaning** | | 2,170 | 1,966 | 1,803 | 1,412 | 7,351 |
| **Unique avian relatives across Mammal-Avian pairings** | | 914 | 1,307 | 1,245 | 1,465 | 4,922 |

**Table S2.** Mutations selected to be introduced into H5N1 backbone for experimental validation

| Segment | Mutation | Count | Freq. % in avian sequences | Freq. % in mammal sequences | Independent mutation occurrence | Species count | Species diversity | Subtype n | Subtype(s) | Notes |
|---|---|---|---|---|---|---|---|---|---|---|
| NP | N21D | 6 | 0.13 | 64.17 | 2 | 3 | swine, felis, human | 4 | H1N1, H1N2, H3N2, H9N2 | Seen in a much higher proportion of mammalian sequences and independently emerges |
| NP | A22T | 57 | 0.85 | 13.22 | 13 | 4 | swine, felis, human, canine | 6 | H1N1, H3N2, H1N2, H7N9, H9N2, H5N1 | Occurs frequently and independently in a range of species |
| NP | V33I | 33 | 5.14 | 65.21 | 10 | 4 | swine, sea lion, human, canine | 6 | H1N1, H1N2, H5N1, H7N9, H9N2, H3N2 | Occurs frequently, independently, seen in various species, high proportion in mammals vs avian |
| NP | R38K | 14 | 0.39 | 21.24 | 6 | 4 | swine, human, canine, skunk | 6 | H1N1, H1N2, H7N4, H7N9, H3N2, H5N1 | Occurs frequently and seen in high proportion in mammals vs avian |
| NP | K48Q | 68 | 0.61 | 14.16 | 2 | 2 | swine, mammal | 3 | H1N1, H3N2, H1N2 | Known mutation |
| NP | R98K | 46 | 1.53 | 15.6 | 9 | 2 | swine, human | 7 | H1N1, H3N2, H1N2, H10N5, H2N2, H7N9, H5N1 | Known mutation |
| NP | V100I | 36 | 0.04 | 27.22 | 5 | 1 | swine | 3 | H3N2, H1N1, H1N2 | Positive control |
| NP | L136I | 7 | 1.22 | 64.58 | 1 | 3 | swine, human, felis | 4 | H1N1, H1N2, H9N2, H3N2 | Seen in a much higher proportion of mammalian sequences |

| | | | | | | | | | | |
|---|---|---|---|---|---|---|---|---|---|---|
| NP | I217V | 10 | 7.65 | 41.75 | 6 | 2 | swine, human | 3 | H1N1, H3N2, H9N2 | Occurs frequently and seen in high proportion in mammals vs avian |
| NP | L283P | 6 | 0.03 | 1.37 | 1 | 1 | human | 2 | H2N2, H1N2 | Positive control |
| NP | A284I | 27 | 0.01 | 10.14 | 1 | 1 | swine | 3 | H1N1, H3N2, H1N2 | Seen in swine, high proportion in mammal vs avian sequences |
| NP | Y289H | 27 | 0.2 | 63.76 | 6 | 3 | swine, human, seal | 4 | H1N2, H1N1, H5N1, H10N7 | Known mutation |
| NP | F313V | | | | | | | | | Not in filtered data but included as a positive control |
| NP | T350K | 15 | 0.13 | 63.82 | 4 | 3 | swine, human, felis | 6 | H1N1, H1N2, H9N2, H3N2, H5N1, H7N9 | Occurs frequently and seen in high proportion in mammals vs avian |
| NP | R351K | 54 | 2.26 | 82.53 | 11 | 3 | swine, human, felis | 6 | H1N1, H3N2, H1N2, H9N2, H7N9, H5N1 | Known mutation |
| NP | Q357K | 33 | 0.21 | 68.73 | 2 | 3 | swine, human, felis | 4 | H1N1, H3N2, H1N2, H9N2 | Positive control |
| NP | R384K | 46 | 4.66 | 17.23 | 10 | 5 | swine, mammal, human, fox, canine | 5 | H1N1, H3N2, H1N2, H7N9, H5N1 | Occurs frequently and independently in a range of species |
| NP | R400K | 8 | 1.75 | 55.8 | 4 | 2 | swine, human | 3 | H1N1, H9N2, H5N1 | Occurs frequently and seen in high proportion in mammals vs avian |
| NP | L418I | 35 | 0.04 | 2.16 | 4 | 4 | felis, canine, cat, polecat | 3 | H3N2, H3N6, H5N1 | Occurs often, in a range of interesting species |
| NP | I425V | 13 | 0.35 | 62.55 | 6 | 3 | swine, human, seal | 4 | H1N1, H1N2, H7N9, H5N1 | Seen in a much higher proportion of mammalian sequences |

| | | | | | | | | | | |
|---|---|---|---|---|---|---|---|---|---|---|
| NP | M426L | 6 | 0.04 | 15.95 | 4 | 2 | swine, human | 5 | H1N1, H1N2, H9N2, H7N9, H3N2 | Seen in a much higher proportion of mammalian sequences |
| NP | R452K | 61 | 13.91 | 65.27 | 12 | 4 | swine, mammal, seal, human | 7 | H1N2, H1N1, H3N2, H5N8, H5N1, H7N9, H9N2 | Occurs frequently and seen in various species, high proportion in mammals vs avian |
| NP | V456L | 51 | 0.28 | 64.29 | 4 | 4 | mink, swine, mammal, human | 6 | H10N4, H3N2, H1N1, H1N2, H7N9, H5N1 | Occurs often, in a range of interesting species, high proportion in mammals vs avian |
| NP | G485R | 20 | 0.03 | 0.2 | 12 | 6 | fox, swine, raccoon, human, seal, mink | 3 | H5N1, H9N2, H7N9 | Occurs frequently and independently in a range of species, yet a rare mutation overall |
| PA | P28S | 34 | 0.1 | 26.9 | 6 | 2 | human, swine | 4 | H1N2, H3N2, H1N1, H7N9 | Occurs frequently and seen in high proportion in mammals vs avian |
| PA | D55N | 19 | 0.9 | 8 | 6 | 4 | human, feline, swine, equine | 6 | H5N1, H7N2, H3N2, H1N1, H9N2, H3N8 | Occurs frequently, independently, and seen in various species |
| PA | R57Q | 55 | 1.75 | 29.74 | 14 | 5 | feline, sea lion, swine, human, mouse | 9 | H7N2, H5N1, H3N2, Undefined, H1N2, H1N1, H7N9, H9N2, H2N2 | Occurs frequently, independently, and seen in various species |
| PA | I62V | 5 | 1.68 | 6.58 | 5 | 5 | skunk, mammal, human, cat, mouse | 4 | H5N1, H1N1, H9N2, H5N6 | Occurs frequently and independently in a range of species, yet a rare mutation overall |
| PA | T85I | 4 | 0.13 | 27.42 | 4 | 1 | human | 3 | H7N9, H9N2, H5N6 | Known mutation |
| PA | T208K | 6 | 0.67 | 17.21 | 4 | 2 | swine, human | 4 | H3N2, H1N1, H1N2, H5N1 | Occurs frequently and seen in high proportion in mammals vs avian |

| | | | | | | | | | | |
|---|---|---|---|---|---|---|---|---|---|---|
| PA | C241Y | 41 | 0.58 | 9.5 | 11 | 5 | mink, skunk, human, canine, swine | 4 | H5N1, H3N2, H1N1, H7N9 | Known mutation |
| PA | N254S | 33 | 0.59 | 30.6 | 5 | 2 | human, swine | 4 | H1N2, H3N2, H1N1, H5N1 | Occurs frequently and seen in high proportion in mammals vs avian |
| PA | R256K | 35 | 0.75 | 32.32 | 9 | 4 | human, canine, swine, ferret | 7 | H5N1, H7N9, H1N2, H3N2, H9N2, H2N2, H1N1 | Occurs frequently, independently, and seen in various species |
| PA | T263E | | | | | | | | | Not in filtered data but included as a high proportion of this mutation in swine |
| PA | P275L | 12 | 1.89 | 30.26 | 8 | 2 | swine, human | 4 | H5N1, H3N2, H7N9, H5N6 | Occurs frequently and independently yet a rare mutation overall |
| PA | S277F | 11 | 1.04 | 2.29 | 6 | 3 | fox, swine, human | 4 | H5N1, H3N2, H7N9, H5N6 | Occurs frequently and independently yet a rare mutation overall |
| PA | K356R | 16 | 12.13 | 39.89 | 7 | 3 | human, swine, mouse | 6 | H5N1, H3N2, H7N9, H2N2, H9N2, H1N1 | Known mutation |
| PA | K362R | 39 | 0.31 | 54.93 | 5 | 3 | pika, human, swine | 5 | H5N1, H1N2, H3N2, H1N1, H7N9 | Known mutation |
| PA | E382D | 14 | 5.64 | 65.77 | 4 | 4 | human, canine, swine, ferret | 5 | H5N1, H1N2, H3N2, H9N2, H1N1 | Occurs frequently, independently, and seen in various species |
| PA | E399K | 25 | 0.92 | 12.85 | 6 | 5 | skunk, canine, swine, mammal, human | 4 | H5N1, H3N2, H1N1, H7N9 | Known mutation |

| | | | | | | | | | | |
|---|---|---|---|---|---|---|---|---|---|---|
| PA | L403I | 22 | 1.11 | 19.58 | 6 | 2 | human, swine | 5 | H9N2, H3N2, H1N2, H1N1, H7N9 | Occurs frequently and seen in high proportion in mammals vs avian |
| PA | D479E | 8 | 0.4 | 3.61 | 6 | 5 | human, equine, canine, mink, fox | 3 | H5N1, H9N2, H7N9 | Occurs frequently and independently in a range of species, yet a rare mutation overall |
| PA | K497R | 26 | 1.03 | 23.78 | 6 | 2 | human, swine | 5 | H5N1, H1N2, H3N2, H1N1, H7N9 | Known mutation |
| PA | I505V | 20 | 2.01 | 21.26 | 10 | 4 | fox, human, swine, mink | 6 | H5N1, H3N2, H1N1, H7N9, H9N2, H5N6 | Occurs frequently, independently, and seen in various species |
| PA | A553S | 4 | 0.43 | 15.35 | 4 | 3 | human, fox, swine | 3 | H9N2, H5N1, H1N1 | Occurs frequently and seen in high proportion in mammals vs avian |
| PA | V602I | 13 | 1.18 | 5.01 | 10 | 4 | human, canine, swine, mink | 6 | H5N1, H5N6, H1N2, H1N1, H7N9, H9N2 | Occurs frequently, independently, and seen in various species |
| PA | K626R | 30 | 10.03 | 30.14 | 7 | 3 | swine, human, cat | 4 | H1N1, H5N1, H7N9, H5N6 | Occurs frequently and seen in high proportion in mammals vs avian |
| PB1 | V12I | 63 | 1.8 | 19.63 | 6 | 3 | human, swine, mammal | 4 | H5N1, H3N2, H1N1, H1N2 | Occurs frequently and seen in high proportion in mammals vs avian |
| PB1 | M40I | 7 | 0.22 | 3.72 | 6 | 4 | sea lion, swine, canine, human | 4 | H5N1, H1N1, H3N2, H1N2 | Occurs frequently and independently yet a rare mutation overall |
| PB1 | N105S | 13 | 0.38 | 0.35 | 5 | 1 | human | 2 | H7N9, H9N2 | Known mutation |
| PB1 | M111I | 29 | 1.42 | 10.35 | 11 | 3 | mink, swine, human | 5 | H5N1, H1N2, H3N2, H1N1, H7N9 | Occurs frequently, independently, and seen in various species, high |

| | | | | | | | | | | |
|---|---|---|---|---|---|---|---|---|---|---|
| | | | | | | | | | | proportion in mammals vs avian |
| PB1 | G154S | 19 | 0.86 | 11.53 | 11 | 5 | feline, mink, human, seal, swine | 6 | H5N6, H10N7, H7N9, H5N1, H1N1, H1N2 | Occurs frequently, independently, and seen in various species, high proportion in mammals vs avian |
| PB1 | M179I | 18 | 4.88 | 59.2 | 8 | 3 | swine, canine, human | 5 | H1N1, H3N2, H7N9, H5N1, H1N2 | Occurs frequently, independently, and seen in various species, high proportion in mammals vs avian |
| PB1 | R211K | 34 | 1.52 | 21.74 | 17 | 4 | seal, cat, swine, human | 8 | H5N1, H1N2, H3N2, H1N1, H7N9, H9N2, H3N8, H2N2 | Occurs frequently, independently, and seen in various species, high proportion in mammals vs avian |
| PB1 | M317I | 34 | 2.92 | 6.99 | 16 | 4 | fox, swine, human, equine | 7 | H5N1, H1N1, H1N2, H3N2, H7N9, H2N2, H3N8 | Known mutation |
| PB1 | S361R | 12 | 0.11 | 54.14 | 5 | 2 | human, swine | 5 | H5N1, H3N2, H2N2, H1N2, H1N1 | Occurs frequently, high proportion in mammals vs avian |
| PB1 | A374S | 14 | 0.52 | 26.98 | 6 | 2 | human, swine | 5 | H7N9, H9N2, H1N2, H3N2, H1N1 | Occurs frequently, independently, high proportion in mammals vs avian |
| PB1 | K433R | 8 | 0.5 | 43.24 | 5 | 2 | human, swine | 4 | H5N1, H3N2, H1N1, H1N2 | Occurs frequently, high proportion in mammals vs avian |
| PB1 | T469I | 23 | 0.3 | 3.49 | 8 | 6 | fox, swine, canine, | 4 | H5N1, H1N1, H3N2, H9N2 | Occurs frequently, independently, and seen in various species |

| | | | | | | | | | | |
|---|---|---|---|---|---|---|---|---|---|---|
| | | | | | | | feline, human, felis | | | |
| PB1 | R486K | 12 | 0.37 | 64.42 | 3 | 1 | swine | 3 | H1N2, H1N1, H3N2 | Occurs frequently, high proportion in mammals vs avian. Swine associated |
| PB1 | S524G | | | | | | | | | Not in filtered data but included as a positive control |
| PB1 | R571K | 26 | 0.61 | 13.06 | 14 | 3 | seal, swine, human | 7 | H10N7, H1N1, H3N2, H7N9, H9N2, H5N1, H1N2 | Occurs frequently, independently, and seen in various species, high proportion in mammals vs avian |
| PB1 | E581D | 8 | 1.48 | 63.01 | 4 | 2 | human, swine | 6 | H9N2, H7N9, H5N1, H1N2, H3N2, H1N1 | Occurs frequently, high proportion in mammals vs avian |
| PB1 | R584Q | 7 | 0.11 | 62.78 | 3 | 2 | swine, equine | 4 | H1N2, H3N8, H3N2, H1N1 | Occurs frequently, high proportion in mammals vs avian |
| PB1 | L598P | 5 | 0.04 | 0.09 | 4 | 1 | human | 2 | H7N9, H5N1 | Known mutation |
| PB1 | Q621R | 40 | 2.27 | 78.48 | 7 | 3 | swine, canine, human | 4 | H1N2, H3N2, H7N9, H1N1 | Known mutation |
| PB1 | S633N | 34 | 0.75 | 14.18 | 17 | 5 | swine, human, raccoon, mouse, mink | 7 | H1N1, H1N2, H9N2, H7N9, H5N1, H3N2, H2N2 | Occurs frequently, independently, and seen in various species, high proportion in mammals vs avian |
| PB1 | S642N | 7 | 96.14 | 46.48 | 6 | 2 | human, swine | 5 | H5N6, H7N9, H9N2, H7N3, H3N2 | Mutation reversal in experimental reference sequence. N642S linked to oselatmivir resistance. |

| | | | | | | | | | | |
|---|---|---|---|---|---|---|---|---|---|---|
| PB1 | M646V | 14 | 0.12 | 1.48 | 7 | 3 | seal, human, swine | 4 | H5N1, H7N9, H9N2, H1N1 | Occurs frequently and independently yet a rare mutation overall |
| PB1 | P708S | 7 | 0.01 | 0.06 | 5 | 1 | human | 2 | H7N9, H5N1 | Known mutation |
| PB1 | I728V | 79 | 1.14 | 22.12 | 3 | 3 | human, swine, mammal | 4 | H7N9, H1N1, H1N2, H3N2 | Occurs frequently and seen in high proportion in mammals vs avian |
| PB2 | K61R | 18 | 2.01 | 34.82 | 9 | 3 | mammal, swine, human | 5 | H1N1, H1N2, H3N2, H7N9, H9N2 | Occurs frequently and seen in high proportion in mammals vs avian |
| PB2 | E65D | 37 | 3.98 | 60.96 | 4 | 2 | human, swine | 5 | H5N1, H1N1, H1N2, H3N2, Undefined | Occurs frequently and seen in high proportion in mammals vs avian |
| PB2 | T76I | 11 | 3.82 | 10.72 | 7 | 4 | felis, equine, swine, human | 5 | H3N2, H3N8, H1N1, H1N2, H7N9 | Occurs frequently and independently in a range of species, yet a rare mutation overall |
| PB2 | N82T | 14 | 0.28 | 5.77 | 8 | 4 | human, swine, cat, caracal | 5 | H5N1, H3N2, H1N1, H1N2, H9N2 | Occurs frequently and independently in a range of species, yet a rare mutation overall |
| PB2 | H127N | 11 | 1.34 | 3.38 | 8 | 3 | human, swine, sea lion | 4 | H5N1, H1N1, H9N2, H7N9 | Occurs frequently and independently in a range of species, yet a rare mutation overall |
| PB2 | I147T | 39 | 9.51 | 66.13 | 14 | 3 | human, swine, caracal | 7 | H5N1, H3N2, H1N2, H1N1, H7N9, H9N2, Undefined | Occurs frequently and seen in high proportion in mammals vs avian |
| PB2 | K157R | 7 | 0.22 | 10.59 | 4 | 2 | swine, human | 3 | H1N1, H5N6, H3N2 | Seen in a much higher proportion of mammalian sequences |

| | | | | | | | | | | |
|---|---|---|---|---|---|---|---|---|---|---|
| PB2 | T184A | 24 | 2.01 | 24.85 | 13 | 4 | human, felis, swine, skunk | 4 | H5N1, H3N2, H9N2, H1N1 | Occurs frequently, independently, and seen in various species |
| PB2 | S225G | 76 | 0.26 | 29.22 | 7 | 6 | swine, felis, mink, panda, equine, human | 6 | H5N6, H3N2, H1N1, H1N2, H3N8, H9N2 | Occurs very frequently, independently, and seen in various species |
| PB2 | T238A | 1 | 0.02 | 1.1 | 1 | 1 | sea lion | 1 | H5N1 | Seen in Sea Lions, rare mutation yet more common in mammals vs avian |
| PB2 | T271A | 36 | 0.26 | 64.32 | 12 | 4 | human, swine, fox, mink | 7 | H5N1, H3N2, H1N2, H1N1, H7N9, H5N6, Undefined | Known mutation |
| PB2 | S286G | 5 | 0.2 | 11.05 | 4 | 1 | human | 3 | H5N1, H10N8, H7N9 | Known mutation |
| PB2 | R299K | 24 | 3.54 | 21.97 | 14 | 3 | human, swine, felis | 8 | H5N1, H1N2, H3N2, Undefined, H9N2, H7N9, H3N8, H1N1 | Occurs frequently, independently, high proportion in mammals vs avian |
| PB2 | R340K | 64 | 28.91 | 71.37 | 14 | 4 | swine, human, mink, equine | 8 | H1N1, H5N6, H3N2, H1N2, H3N8, H7N9, H9N2, Undefined | Occurs very frequently, independently, and seen in various species |
| PB2 | E391D | 11 | 0.38 | 30.91 | 6 | 1 | swine | 4 | H3N2, H5N1, H1N2, H1N1 | Occurs frequently, high proportion in mammals vs avian. Swine associated |
| PB2 | V480I | 26 | 2.88 | 13.08 | 16 | 3 | human, swine, raccoon | 5 | H5N1, H3N2, H9N2, H1N2, H1N1 | Occurs frequently, independently, and seen in various species |
| PB2 | V560L | 45 | 0.16 | 40.92 | 4 | 1 | swine | 4 | H1N1, H1N2, H3N2, Undefined | Occurs very frequently and seen in high proportion in mammals vs avian |

| | | | | | | | | | | |
|---|---|---|---|---|---|---|---|---|---|---|
| PB2 | K574R | 17 | 0.66 | 4.39 | 10 | 4 | human, swine, mammal, skunk | 6 | H5N1, H3N2, H1N1, H1N2, H9N2, H5N6 | Occurs frequently and independently in a range of species, yet a rare mutation overall |
| PB2 | V584I | 35 | 2.92 | 11.06 | 9 | 2 | swine, human | 5 | H9N2, H1N2, H3N2, H1N1, H7N9 | Occurs frequently and independently, yet a rare mutation overall |
| PB2 | G590S | 18 | 4.19 | 64 | 6 | 3 | human, equine, swine | 7 | H5N1, H3N8, H7N9, H1N1, H3N2, Undefined, H1N2 | Known mutation |
| PB2 | Q591R | 35 | 0.14 | 62.86 | 8 | 2 | human, swine | 6 | H5N1, H3N2, H7N9, H1N1, Undefined, H1N2 | Known mutation |
| PB2 | E627K | 426 | 1.09 | 10.09 | 143 | 13 | human, fox, otter, canine, mustela, mink, lynx, swine, seal, mouse, bear, raccoon, skunk | 10 | H5N6, H5N1, H7N4, H9N2, H7N7, H10N8, H7N9, H3N8, H5N8, H1N1 | Positive control |
| PB2 | M645L | 8 | 0.45 | 72.51 | 2 | 2 | felis, swine | 4 | H3N2, H1N1, Undefined, H1N2 | Seen in a much higher proportion of mammalian sequences |
| PB2 | D701N | 149 | 0.09 | 17.19 | 56 | 14 | human, equine, tiger, mink, lynx, swine, mouse, fox, seal, skunk, mustela, sea | 12 | H5N6, H5N1, H3N8, H7N7, H9N2, H7N9, H5N8, H1N2, H10N5, H10N7, H1N1, H3N2 | Positive control |

| | | | | | | | |
|---|---|---|---|---|---|---|---|
| | | | | | | | lion, feline, bear |

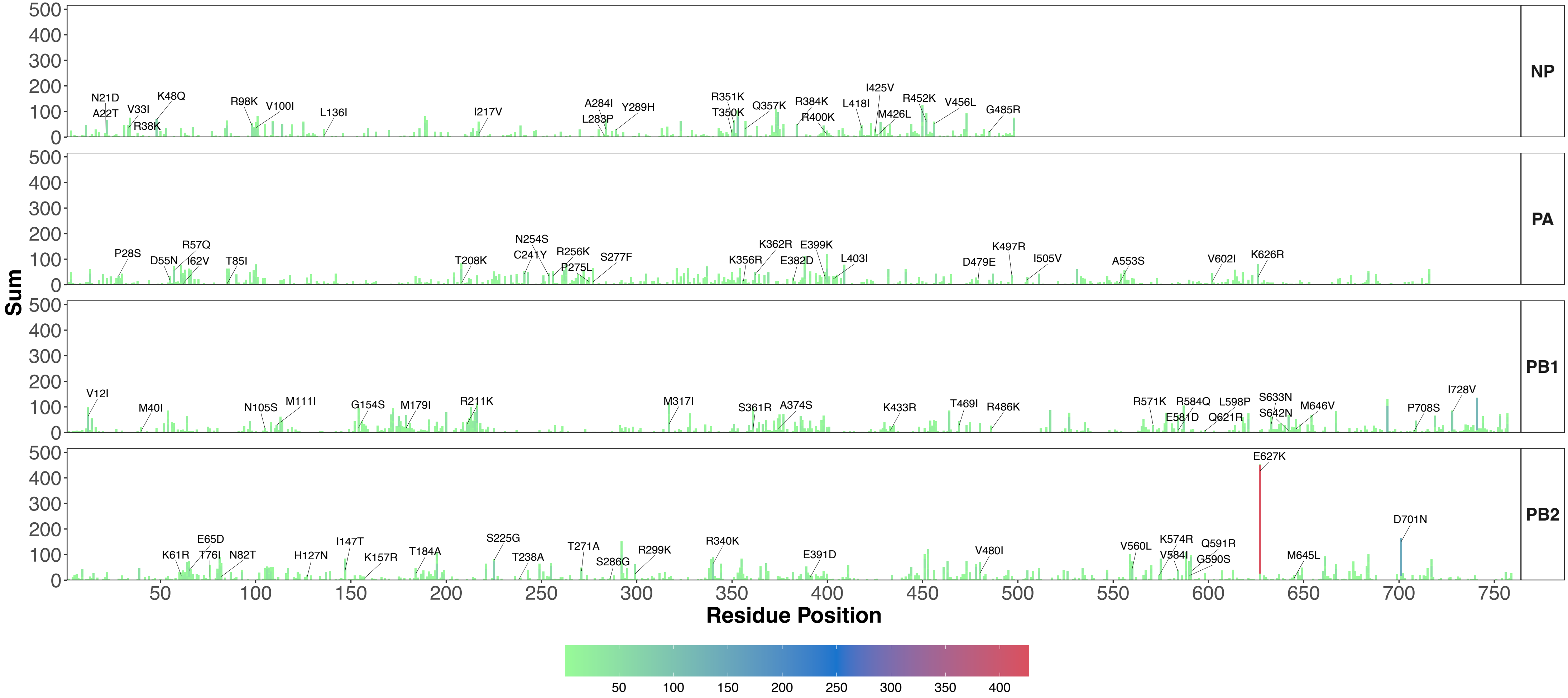

**Figure S1**. Sum of all mutations at residue stacked by individual mutation for each polymerase segment and NP. Mutations selected for experimental analysis labelled.

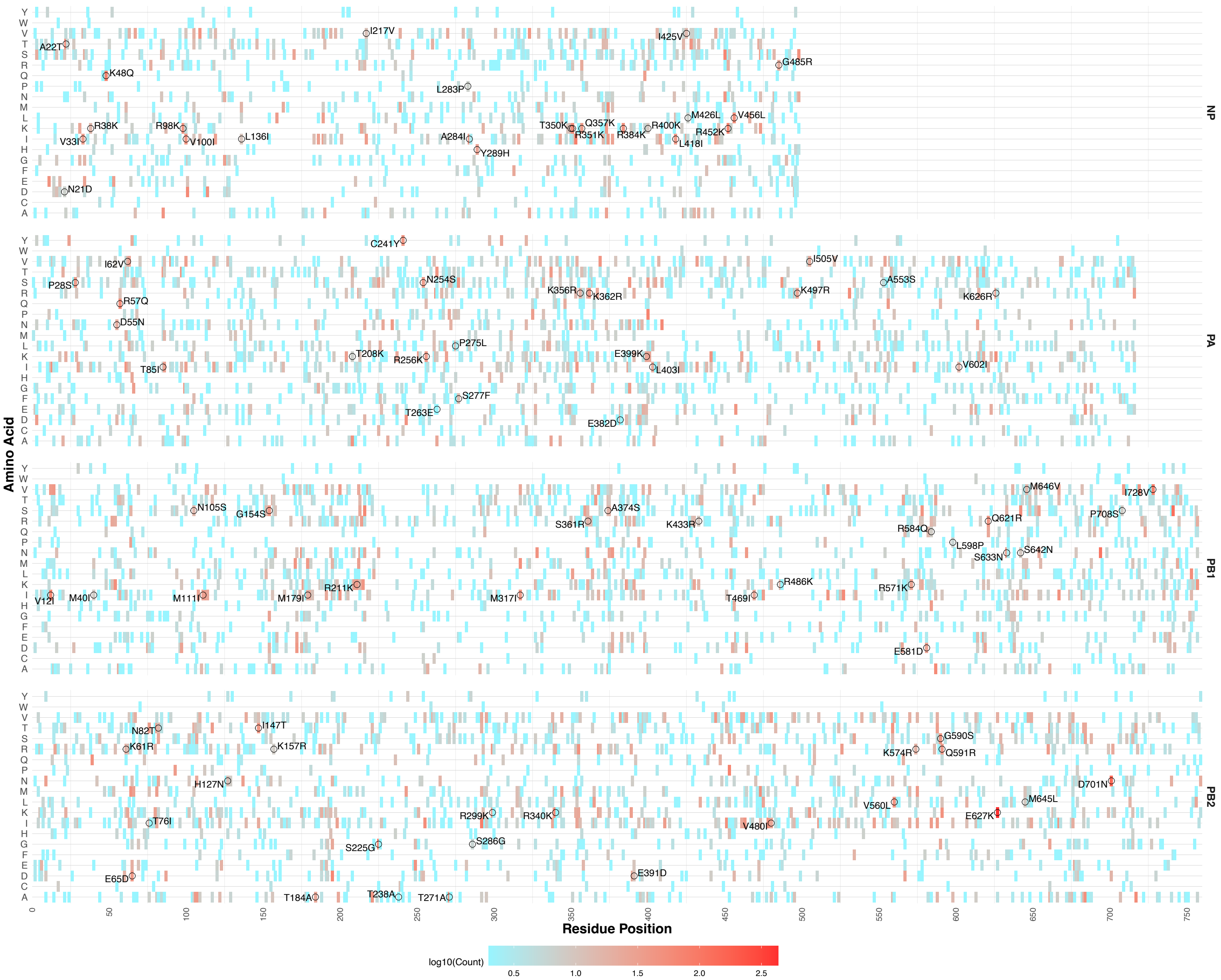

**Figure S2.** Heatmap of amino acid residues seen in mammalian sequences. Coloured by frequency on a log10 scale. Mutations selected for analysis circled and labelled.

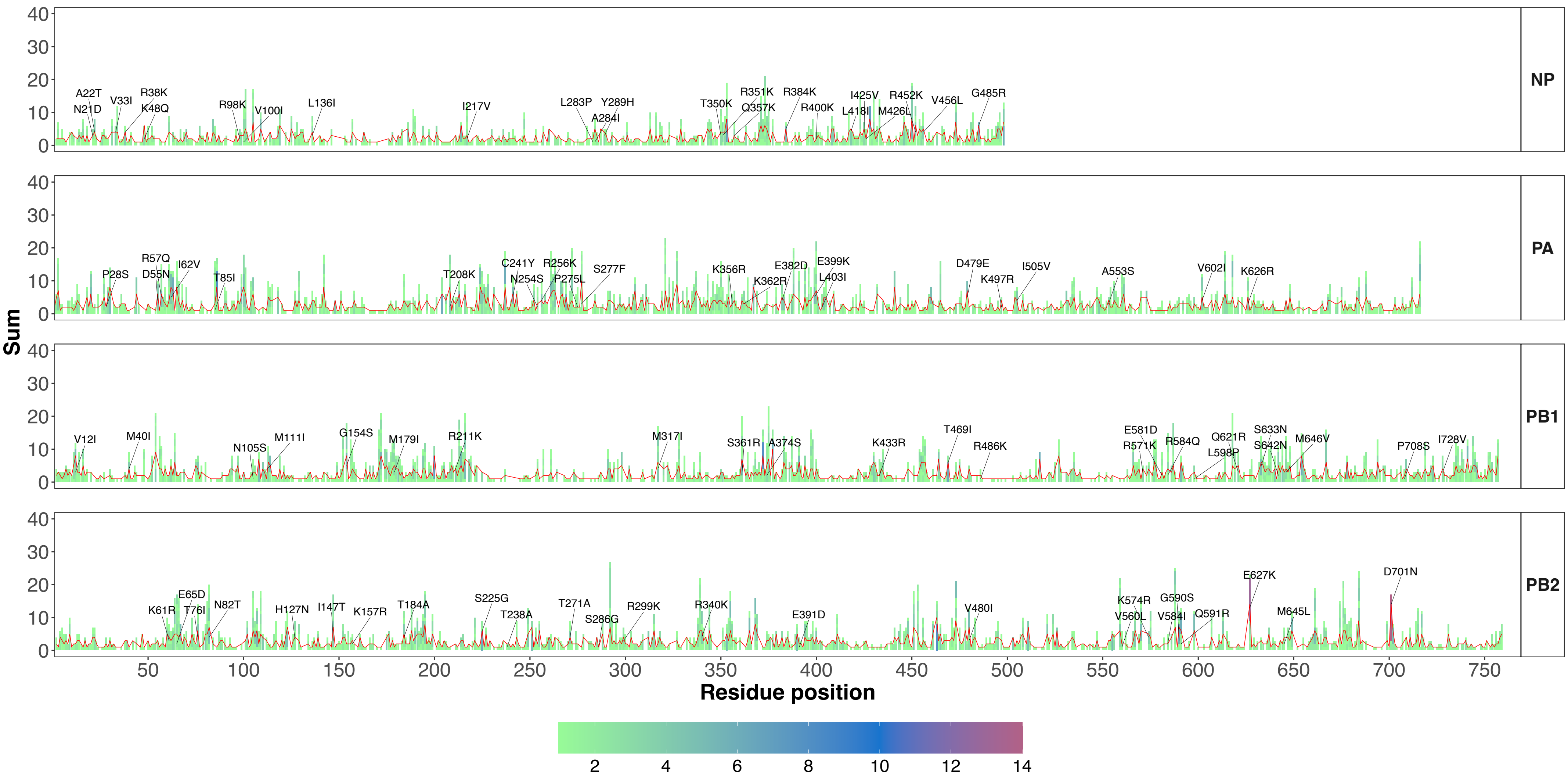

**Figure S3.**

Stacked count of unique species associated per mutation. Red line overlaid displays unique species at residue position. Mutations selected for experimental analysis labelled.

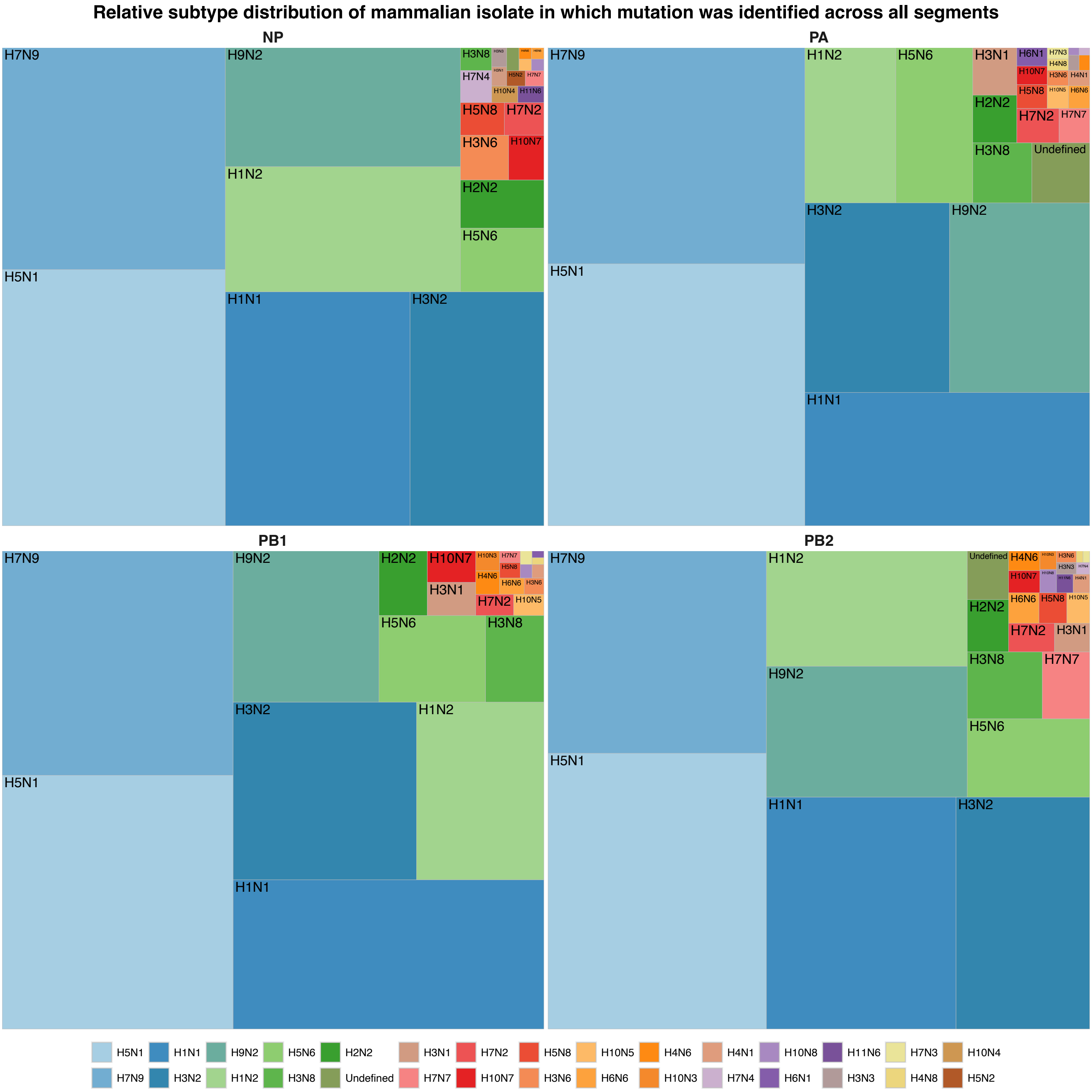

**Figure S4.**
Distribution of subtypes based on mammalian isolate from which mutations were identified by polymerase segment (PA, PB1, PB2) and NP.

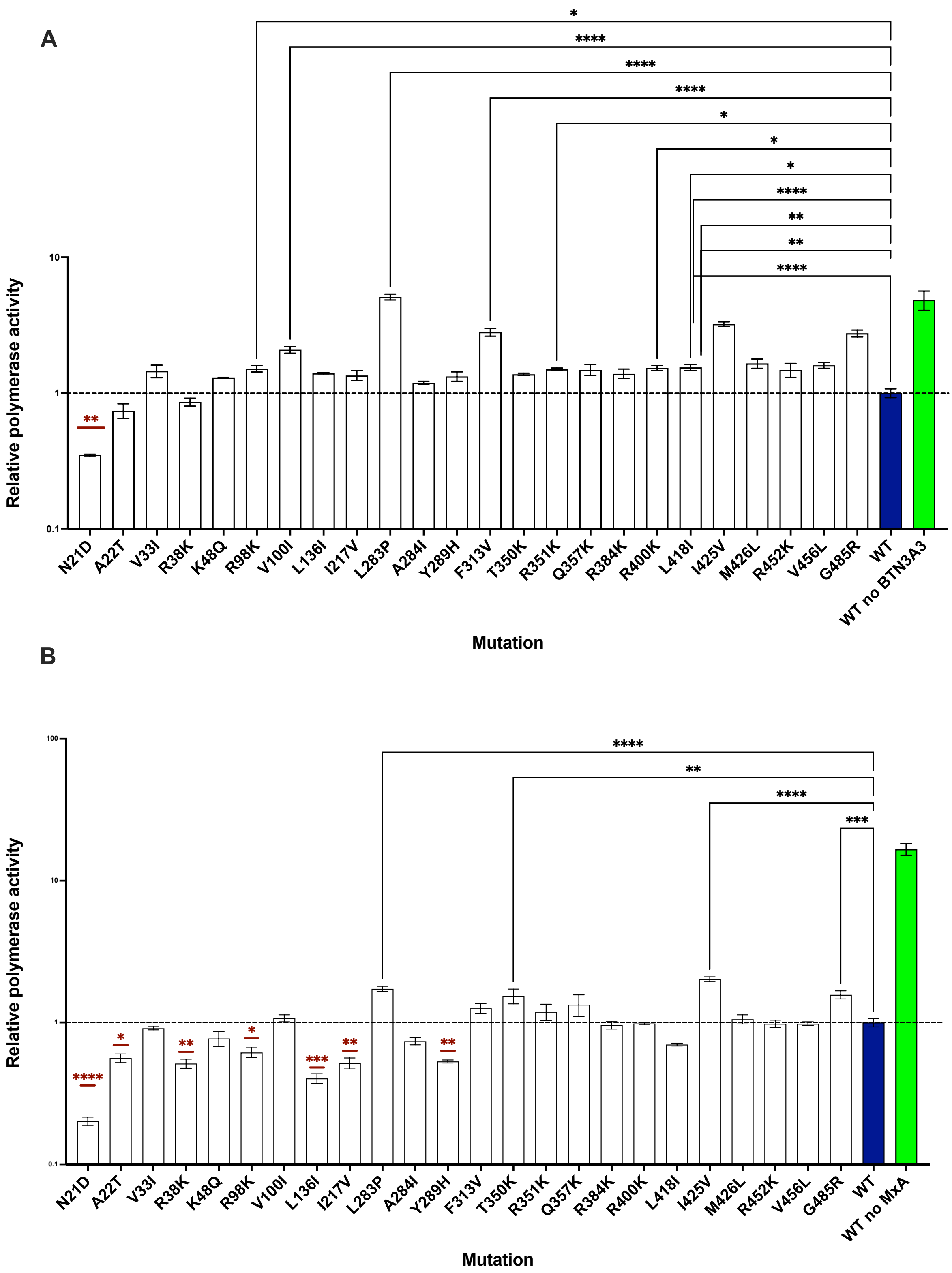

**Figure S5.**

Minigenome assay of NP mutations in human 293T cells with restriction factor Mx1 **(A)** or BTN3A3 **(B)** exogenously expressed. Polymerase activity displayed as Firefly/Renilla normalised to wild type, with SEM bars. Results of multiple comparisons one-way ANOVA comparing WT to mutant are displayed. (*P ≤ 0.05, ** P ≤ 0.01, *** P ≤ 0.001, **** P ≤ 0.0001). Results from a representative minigenome are shown. WT with no restriction factor shown in green for reference but not included in statistical analysis.

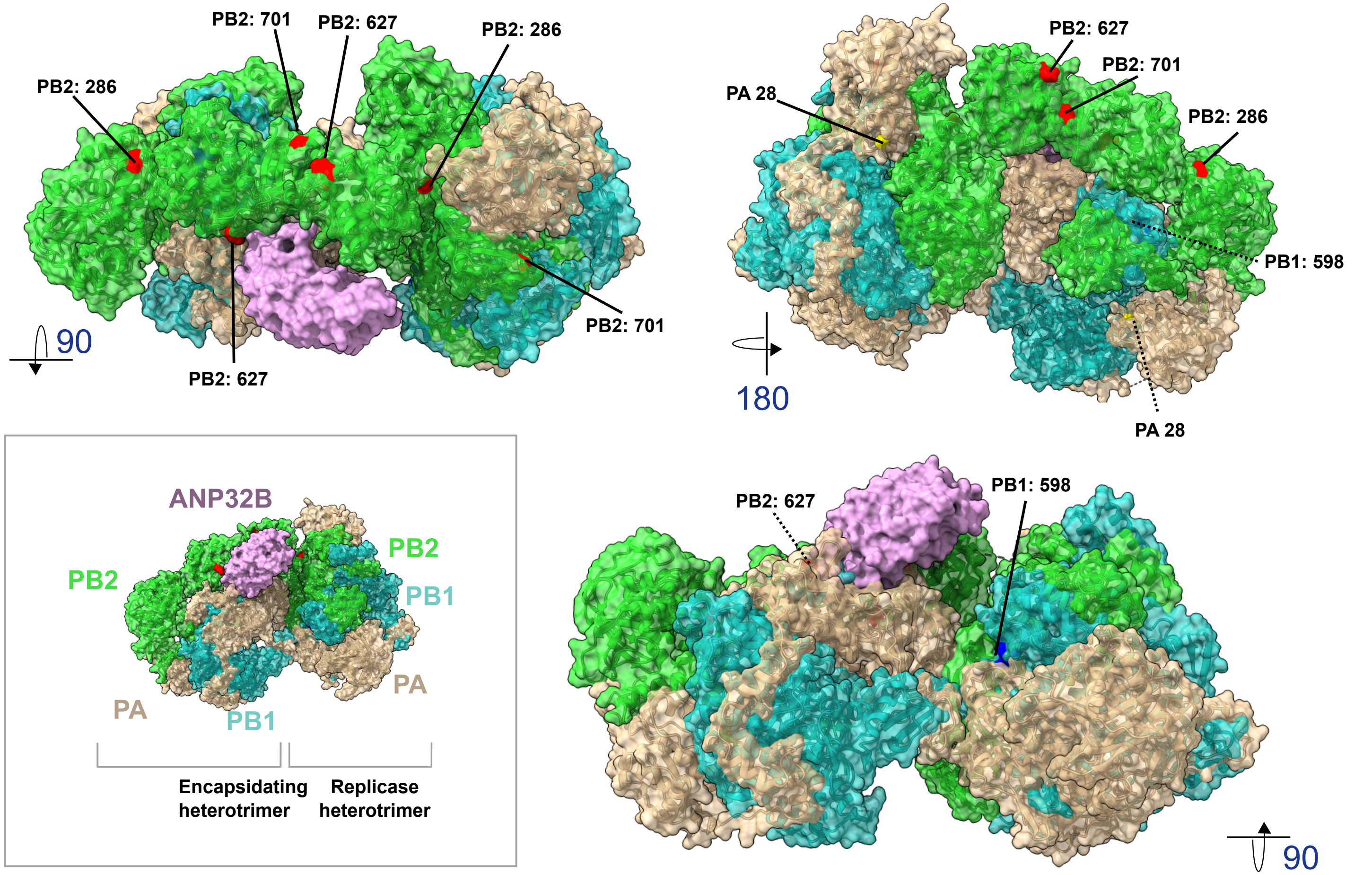

**Figure S6:**

H5N1 asymmetric polymerase dimer bound to human ANP32B (PDB: 8R1J). Dessert: PA. Green: PB2, Blue: PB1, Purple: ANP32B. PB2 627, 701, 286, PA 28 and PB1 598 labelled. PB2 627 and 701 are in the ANP32 binding interface in the polymerase asymmetric dimer configuration, however PB2 286 and PB1 598 are not.

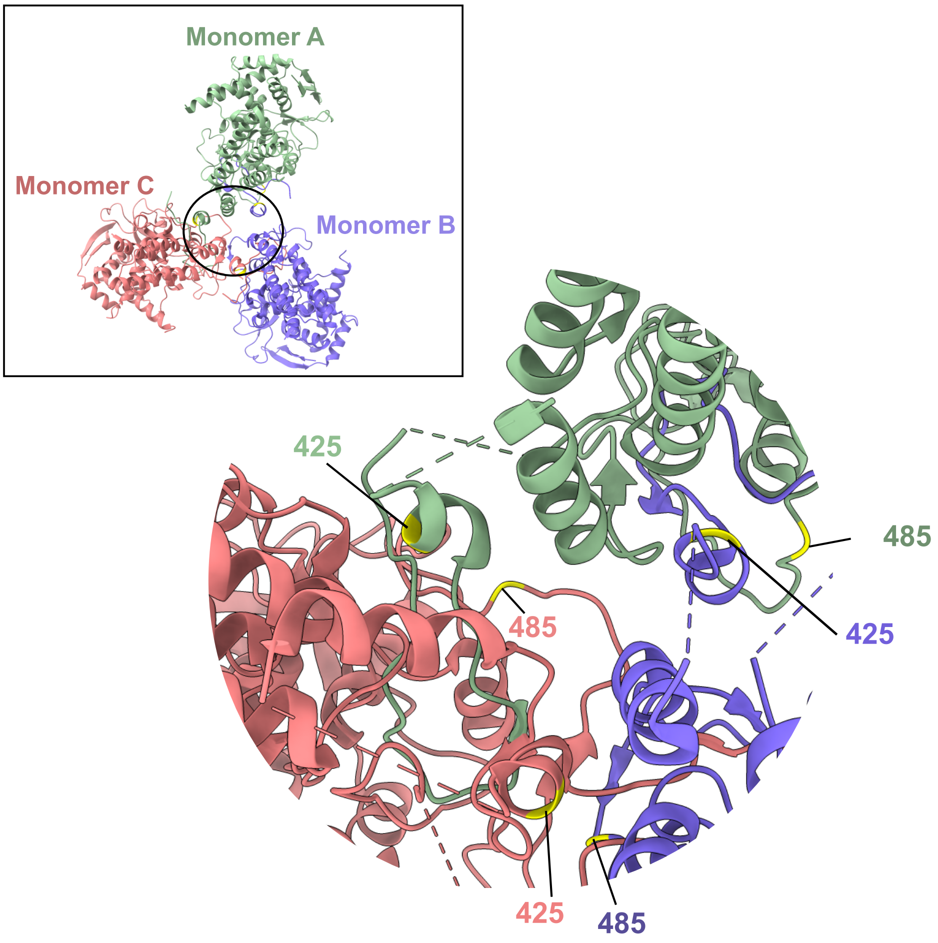

**Figure S7:**
H1N1 NP trimer structure (PDB:2Q06) with NP 485 and 425 labelled. Residue 425 is located in the tail loop and 485 is in the main body. Interactions with the tail loop and body domain occur during oligomerisation when NP monomers interact forming a trimer structure.
